# Transcriptional Activation of Regenerative Hematopoiesis via Vascular Niche Sensing

**DOI:** 10.1101/2023.03.27.534417

**Authors:** Tomer Itkin, Sean Houghton, Ryan Schreiner, Yang Lin, Chaitanya R. Badwe, Veronique Voisin, Alex Murison, Negar Seyedhassantehrani, Kerstin B. Kaufmann, Laura Garcia-Prat, Gregory T. Booth, Fuqiang Geng, Ying Liu, Jesus M. Gomez-Salinero, Jae-Hung Shieh, David Redmond, Jenny Z. Xiang, Steven Z. Josefowicz, Cole Trapnell, Joel A. Spencer, Lior Zangi, Brandon Hadland, John E. Dick, Stephanie Z. Xie, Shahin Rafii

**Author notes:** Correspondence should be addressed to Tomer Itkin and Shahin Rafii.

## Abstract

Transition between activation and quiescence programs in hematopoietic stem and progenitor cells (HSC/HSPCs) is perceived to be governed intrinsically and by microenvironmental co-adaptation. However, HSC programs dictating both transition and adaptability, remain poorly defined. Single cell multiome analysis divulging differential transcriptional activity between distinct HSPC states, indicated for the exclusive absence of Fli-1 motif from quiescent HSCs. We reveal that Fli-1 activity is essential for HSCs during regenerative hematopoiesis. Fli-1 directs activation programs while manipulating cellular sensory and output machineries, enabling HSPCs co-adoptability with a stimulated vascular niche. During regenerative conditions, Fli-1 presets and enables propagation of niche-derived Notch1 signaling. Constitutively induced Notch1 signaling is sufficient to recuperate functional HSC impairments in the absence of Fli-1. Applying FLI-1 modified-mRNA transduction into lethargic adult human mobilized HSPCs, enables their vigorous niche-mediated expansion along with superior engraftment capacities. Thus, decryption of stem cell activation programs offers valuable insights for immune regenerative medicine.

---

To sustain hematopoiesis, to continuously produce immune blood cells throughout adulthood, HSCs possess a preset molecular epigenetic program that firmly maintains a developmentally predefined path, even under various stress-induced conditions^1–3^. Nevertheless, the multicellular HSC niche landscape, with its distinct and heterogeneous arsenal of cellular components, provides differential instructive cues, essential for maintenance of balanced hematopoiesis^4–6^, even by hematopoietic progeny^7^. Thus, there must be an adaptive two-way crosstalk between HSCs and their dynamic niche cells augmenting and fine-tuning steady state and stress-induced production of immune blood cells while maintaining the HSC pool^8^, lending credence to an undefined intrinsic HSPC transcriptional-mechanism regulating this crosstalk. Notably, the molecular hubs that coordinate sensing and communication of stem cells with their interactive niche cells to define between multiple stem cell fates, remain unknown. Uncovering these hubs and their defined roles may assist translational efforts to manipulate hematopoietic stem cell fate for immune regenerative medical resolutions.

## In depth interrogation of dynamic HSPC states reveals absence of Fli-1 transcriptional activity in non-active quiescent HSCs

In search of transcriptional hubs coordinating the HSC-niche adaptability and crosstalk, we comprehensively probed a murine perivascular niche residing population of HSPCs, highly enriched for long term repopulating (LTR) HSCs^9^ phenotypically marked as CD150^+^/CD48^negative^/Lineage^negative^/Sca-1^+^/c-Kit^+^, SLAM LSK HSPCs. This population of sorted cells was subjected to combined multimodal single nuclei RNA- and ATAC-seq (snRNA/ATAC-seq) analysis to define transcriptional output and chromatin accessibility states (Extended Data Fig. 1a). Annotating cell identity for clusters over weighted nearest neighbor (WNN) uniform manifold approximation and projection (UMAP) analysis, we have identified clusters for hematopoietic stem cells (HSC), hematopoietic multi-potent progenitors (MPP), progenitors for platelet producing megakaryocytes (MkP) and for erythrocytes (EryP), confirming a previous report for SLAM LSK HSPCs^10^ (Extended Data Fig. 1b). To transcriptionally authorize the location and identity of HSC derived nuclei over our WNN UMAP, we applied an HSC transcriptional signature recently defined for engraftable LTR-HSCs, achieved with application of barcoding-based single cell lineage tracing^11^. Notably, although confirming the HSC cluster identity, two distinct centers with the highest Kernel denseties were observed (Extended Data Fig. 1c), suggesting distinguished sub-populations in the HSC cluster. Differential nuclear RNA transcriptome analysis confirmed the identity of each cluster with known gene markers for each HSPC sub-type (Extended Data Fig. 1d **and Supplementary Table 1**). Differential motif activity based on chromatin accessibility analysis further exhibited transcription factor (TF) activity of known HSC factors, in the HSC cluster, among numerous other unique TFs for each cluster that were not defined yet for their role in hematopoiesis (Extended Data Fig. 1e **and Supplementary Table 2**).

To further uncover additional HSC and MPP sub-clustering potential, we re-clustered only the HSC and MPP single nuclei following exclusion of MkP and EryP clusters, and reapplied HSC transcriptional signature to confirm localization of HSC sub-clusters (Fig. 1a). New HSC/MPP clustering WNN UMAP analysis exhibited two HSC clusters (HSC1 and HSC2), which corelate with the two density centers harboring the highest HSC signature score. Two MPP clusters (MPP1 and MPP2), and an HSC-MPP mixed cluster with low HSC signature score were further identified (Fig. 1b). Examining differential transcriptome expression and motif activity in HSC/MPP sub-clusters (**Supplementary Table 3**), we noted uppermost activity of Spi family TFs in the late MPP2 cluster but no activity in the HSC clusters (Fig. 1c). This finding supports the notion that Spi activity in HSCs may be restricted to stress-induced regenerative conditions^12^, but not during steady state. The earlier MPP1 cluster was defined by GATA TF family activity, while the transitioning HSC-MPP cluster was predominantly defined by the accesability of Nfi family of TFs (Fig. 1c). Comparing the gene ontology (GO) enrichment results analysis for biological processes and motif activity for the distinct HSC clusters, we noted that the HSC1 cluster exhibited an active proliferative signature with activity of self-renewal associated TFs, including Rbp_j_^13^ and Stat3/5^14, 15^ (Fig. 1c, d, **and Supplementary Table 3**). While, cluster HSC2 represents the metabolically non-active population of HSCs with active TF motifs maintaining murine stem cell quiescence such as Ctcf^16^ and Hlf^17^, additionally it displayed enrichment for negative regulation of multiple active cellular processes (Fig. 1c, d, **and Supplementary Table 3**). Examining the top enriched active motifs portraying each HSC/MPP subclusters, we noted the unique lack of activity of the hemato-vascular TFs Erg and Fli-1 in the quiescent and non-active HSC2 cluster versus the other clusters (Fig. 1c, d, **and Supplementary Table 3**).

**Fig. 1:**
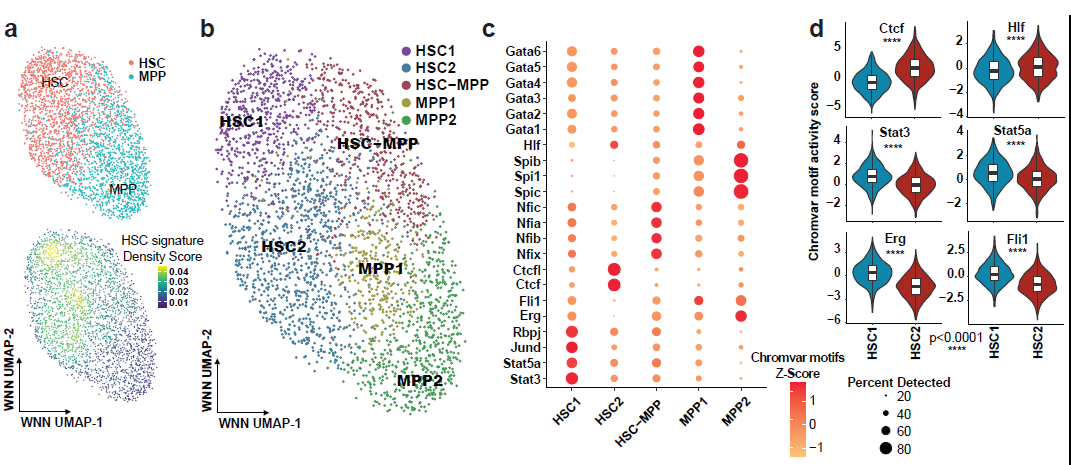
Deep analysis of perivascular HSPCs reveals unique motif activity signatures per transitional state. **a**, WNN UMAP of single nuclei from selected HSC and MPP sub-clusters without the MkP and EryP sub-clusters (upper panel). Engrafting LTR-HSC transcriptional signature (from Rodriguez-Fraiticelli, *et al.* 2020) assigned on a WNN UMAP space for HSC and MPP clusters (lower panel). **b**, WNN UMAP projection of HSC/MPP re-clustering with cell identity annotation. **c**, Dot plot of ChromVAR motif activity of selected motifs in each HSC/MPP sub-cluster. Colors and sizes of dots indicate the level of motif activity by Z-score and percent of cells displaying the activity for the indicated motif, respectively, in each HSC/MPP sub-cluster. **d**, Violin-plots displaying chromVAR motif activity score for selected motifs per HSC sub-clusters HSC1 (blue) and HSC2 (red). Wilcoxon Rank Sum Test was used to determine p-Values.

We have recently found that expression of Erg and Fli-1 in endothelial cells, is essential to uphold vascular transcriptional physiological programs, and thus hypothesized that they may perform similarly for activated HSCs^18^. While the role of Erg in regulation of HSC activity and balancing between self-renewal to differentiation was established^19, 20^, the role of Fli-1 (Friend Leukemia integration-1)^21^ is poorly characterized in HSC regulation.

Global embryonic Fli-1 deficiency results in fetal lethality attributed to vascular leakiness, and adult hematopoietic inducible Fli-1 genetic deletion impairs development and maturation of some hematopoietic myeloid lineages, affecting also the numbers of and skewing between myeloid progenitors^22, 23^, and directs megakaryocyte cell fate specification^24^. Additionally, Fli-1 regulates lymphocyte progeny by restricting formation of memory NK cells^25^ and by restraining effector T cell lineage differentiation^26^. Nonetheless, the physiological role of the Fli-1 TF in modulating active versus quiescent states of adult immune stem cells was never defined.

## Adult hematopoietic Fli-1 deficiency induces thrombocytopenia and impairs HSC immune regenerative capabilities

To formally determine the role of Fli-1 in adult HSCs and its role in the regulation of adult hematopoiesis, we genetically induced Fli-1 deletion in adult mice. Global and hematopoietic specific Fli-1 deficiency in mice results in rapid mortality due to severe thrombocytopenia (Extended Data Fig. 2a-k).

Further examining HSPC status in the bone marrow (BM), we observed failure of committed progenitors to form megakaryocytic colonies, whereas the BM displayed a typical myeloid progenitor “stress response” as evidenced by enhanced numbers of both granulocyte-macrophage colonies and phenotypically defined Lineage^negative^/Sca-1^+^/c-Kit^+^ (LSK) HSPCs (Extended Data Fig. 2l, m). By contrast, we noted a decrease in the number of the more primitive and stem cell enriched SLAM LSK HSPCs (Extended Data Fig. 2n). This potential stem cell impairment was further corroborated by the finding that HSCs within the Fli-1^ROSAΔ^ LSK cells, failed to engraft following transplantation (Extended Data Fig. 2o, p). Thus, Fli-1 deficiency manifests HSC stress-response, repopulation, and engraftment defects.

To exclude niche homing defects of Fli-1^ROSAΔ^ LSK HSPCs, we examined HSPC BM trafficking capacity by intravital microscopy 24 hours following transplantation into normal wild-type irradiated mice. We noted that while a small subset of Fli-1^ROSAΔ^ HSPCs (19%) were wedged at the luminal side of the vessel wall, the majority of Fli-1^ROSAΔ^ HSPCs managed to home into the BM as well as 100% of WT HSPCs (Extended Data Fig. 3a, b **and Supplementary Videos 1, 2**).

Speculating that this small subset of cells failing to home to the BM could represent repopulating HSCs, we transplanted LSK HSPCs prior to Fli-1 deficiency induction and administrated tamoxifen a week later, ensuring proper BM lodgment. Again, Fli-1^ROSAΔ^ HSCs failed to repopulate transplanted mice (Extended Data Fig. 3c, d). Furthermore, mixed competitive transplants with WT cells and induction of Fli-1 deletion 4 months post-transplant revealed a gradual multi-lineage collapse of hematopoiesis derived from Fli-1^ROSAΔ^ HSPCs, with a very rapid deterioration of the myeloid lineage followed by a decline in the lymphoid lineages (Extended Data Fig. 3e-h). Enhanced frequency of apoptotic HSPCs was observed following Fli-1 deficiency induction in this model (Extended Data Fig. 3i). Thus, Fli-1 deficiency in HSCs results in immune lineage developmental defect, impairment of survival, and hampered self-renewal/differentiation capacities, preventing proper immune regeneration and recovery.

## Fli-1 deficient HSPCs transcriptionally transition into a quiescent state

To further asses if and how Fli-1 confers niche-guided regenerative potential to HSPCs, we leveraged our previously described vascular niche-based regenerative HSPC expansion system^27^. This approach mimics physiological stress-induced regenerative conditions and permits expansion of a sufficient pool of HSPCs that can be isolated for multi-omics analyses (Extended Data Fig. 4a). In this system, Fli-1 deficient HSPCs failed to expand in co-culture with vascular niche cells. Notably, Fli-1 deficient HSPCs were not responsive to the microenvironmental signals, as no beneficial effect for Fli-1^ROSAΔ^ HSPCs expansion was observed in the presence of a vascular niche versus no niche in presence (Fig. 2a **and** Extended Data Fig. 4b). Analyzing expansion dynamics in co-culture, we noted hematopoietic arrest and collapse very shortly following induction of Fli-1 deletion (**Supplementary Videos 3, 4**). Therefore, Fli-1 deficiency may result in the impairment of HSPCs to sense incoming activation cues from a prototypical niche, such as a vascular niche.

**Fig. 2:**
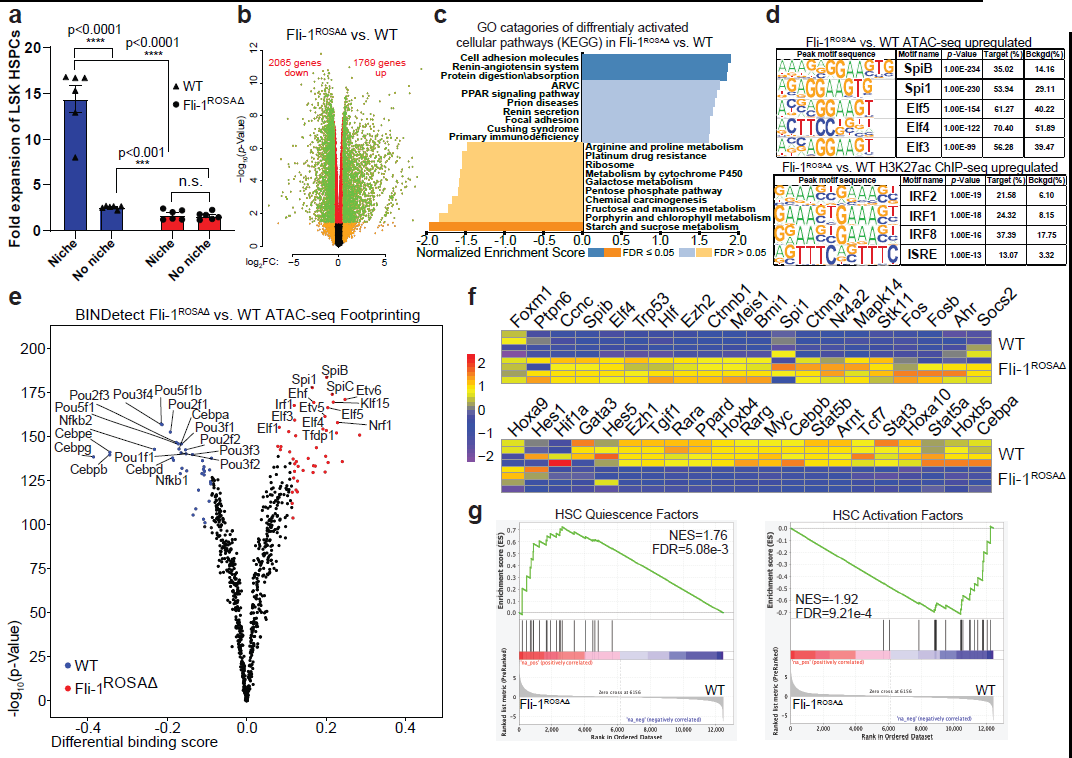
Fli-1 counterbalances niche-dependent HSPC regenerative activation versus conservative quiescence programs. **a,** BM LSK HSPCs were sorted and introduced into a serum free co-culture condition with or without a vascular niche. Frequency of LSK HSPCs was determined by flow cytometry and fold expansion was calculated, relatively to initial number of seeded cells. Two-way ANOVA multiple comparisons was used; n=6 BM donor mice per genotype, each point represents an average of 3 technical replicates. **b,** Volcano plot of differentially expressed genes in Fli-1^ROSAΔ^ vs. WT HSPCs. Thresholds were set at |LogFC|>0.5 for x-axis and at FDR<0.05 for y-axis. Green and red color labeling is for genes with FDR<0.05 and green and yellow color labeling is for genes with |LogFC|>0.5. See also Table S1. **c,** Gene ontology (GO) categories of biological processes enrichment for differentially expressed genes in RNA-seq analyses of Fli-1^ROSAΔ^ vs. WT HSPCs. Positively enrichment scored processes are enriched in Fli-1^ROSAΔ^ HSPCs while negatively enrichment scored processes are enriched in WT HSPCs. **d,** Transcription factor binding site motif enrichment analysis for upregulated ATAC-and H3K27ac ChIP-seq peak sites in Fli-1^ROSAΔ^ vs. WT HSPCs. See also Tables S2 and S3. **e,** Volcano plot of differential BINDetect motif footprinting in Fli-1^ROSAΔ^ vs. WT HSPCs. Blue and red colors are for labeling of footprinting enriched in WT HSPCs (Blue) and in Fli-1^ROSAΔ^ HSPCs (Red), above the set thresholds for the binding score and for the p-Value. **f,** Heatmap for selected HSC quiescence (upper panel) and HSC activation (lower panel) factors across RNA-seq replicates of Fli-1^ROSAΔ^ and WT HSPCs. Scale bar and coloring represent the Z-score scaled by gene for Log counts per 10^6^ normalized by library size (trimmed mean of M-values). **g,** GSEA analysis plot for HSC quiescence (left panel) and HSC activation (right panel) factors expression in Fli-1^ROSAΔ^ vs. WT HSPCs, showing a positive enrichment for quiescence factors and negative enrichment for activation factors in Fli-1^ROSAΔ^ HSPCs.

Differential gene expression analysis, of Fli-1^ROSAΔ^ versus WT HSPCs undergoing expansion in co-culture with a vascular niche, revealed alteration in thousands of genes (Fig. 2b **and Supplementary Table 4**). Gene set enrichment analysis for biological processes displayed enhanced biological activities associated with quiescence among Fli-1^ROSAΔ^ HSPCs. These represented processes involved in cellular quiescence such as adhesion^28^, protein digestion and absorption^29^, alongside with a drop out in activation of related metabolic pathways^30^ (Fig. 2c), which are essential to maintain fit interactions between HSCs with their niche^31^.

Next, we examined differential chromatin landscape accessibility and chromatin modification for transcriptional priming patterns, by ATAC-seq and H3K27ac ChIP-seq analyses, between Fli-1^ROSAΔ^ and WT expanding HSPCs in co-cultures (**Supplementary Tables 5, 6**). Motif enrichment analysis (MEA) for differential ATAC and H3K27ac ChIP peaks enriched in promoter/enhancer regions from Fli-1^ROSAΔ^ HSPCs presented a signature of elements restricting HSC activation, while sustaining and enforcing HSC quiescence, such as the TFs Pu.1 (Spi1)^32^ and IRF2^33^ (Fig. 2d). Moreover, Spi and IRF factors partner physically in HSCs by IRFs binding to Spi’s PEST domain^34^. Notably, our MEA ATAC-seq signature was almost identical to Spi1’s ChIP-seq MEA signature, that was performed on primitive and mostly quiescent HSPCs^12^. Additionally, examining the MEA signature of ATAC and H3K27ac ChIP peaks enriched for WT HSPCs, we confirmed that Fli-1 deletion in HSPCs results with loss of chromatin accessibility and chromatin priming of binding domains related to Fli-1 and other homologous ETS TFs (Extended Data Fig. 4c). Further employing our ATAC-seq data, we performed computational “footprinting” analysis^35^, to reveal TF binding activity. This approach further indicates that Fli-1^ROSAΔ^ HSPCs contain higher predicted activity of Spi family members (Fig. 2e).

Overall, our data qualifies Fli-1 as a transcriptional master regulator in HSPC regenerative activation, possibly counterbalancing Spi1, during regenerative hematopoiesis. To confirm Fli-1’s position as a master TF, switching between quiescence to activation programs, we evaluated changes in expression of HSC TFs known to regulate quiescence or activation. We determined a significant transcriptional induction of quiescence program genes in Fli-1^ROSAΔ^ HSPCs along with a significant transcriptional cellular activation program shutdown in Fli-1^ROSAΔ^ HSPCs (Fig. 2f, g).

## Fli-1 presets niche co-adoptability through Vegfa/Notch signaling

In order to gain mechanistic insight regarding processes which augment Fli-1 dependent hematopoietic activation through interaction with the vascular niche, we queried the sets of genes that were suppressed both on the RNA and chromatin accessibility levels in Fli-1^ROSAΔ^ HSPCs (Extended Data Fig. 5a), presuming that those are controlled by a global Fli-1 targeted program. GO for biological pathways analysis revealed that Fli-1^ROSAΔ^ HSPCs were transcriptionally inferior to respond to various types of external stimuli, mainly immune stress associated inflammatory activation cues, and displayed impaired capacity for cytokine production (Extended Data Fig. 5b), two features which are crucial for the interaction and crosstalk between HSPCs to their neighboring niche cells. To confirm this finding, we examined the transcriptome of HSPC sensory elements, and in Fli-1^ROSAΔ^ HSPCs we distinguished a significant suppression of known hematopoietic extracellular activating sensory components while noting elevated expression in quiescence promoting ones (Extended Data Fig. 5c, d).

This data indicates that a master transcriptional regulator like Fli-1 presets HSPC activation programs and prepares HSCs to prompt into a regeneratively active state through sensing of the relevant augmenting signals arriving from a cytokine-stimulated microenvironment.

We and others have shown that activation of Notch signaling in HSPCs by vascular niche-expressed Notch ligands augments HSPC developmental choices such as regenerative expansion while limiting premature steady state differentiation^27, 36^. This crosstalk with the vascular niche to reciprocally activate Notch signaling is initiated by HSPCs via secretion of Vegfa to stimulate specialized vascular ECs^27, 37^. Thus, dysregulation of Vegf/Notch pathways might explain Fli-1^ROSAΔ^ HSPC insensitivity to vascular niche presence (Fig. 2a) and support the observed suppression of extracellular sensing elements along with cytokine production defects in Fli-1^ROSAΔ^ HSPCs (Extended Data Fig. 5b). To address this hypothesis, we interrogated the regulatory role of Fli-1 in masterminding the reciprocal Vegf/Notch pathways crosstalk in HSPCs. Negative dysregulation for multiple Notch and Vegf pathway elements was observed in Fli-1^ROSAΔ^ HSPCs, among which Vegfa RNA was dramatically downregulated (Fig. 3a, b). Although the requirement for Notch signaling in HSC maintenance is debated^38^, Notch1 activation was shown to augment HSC self-renewal^39, 40^ while active Notch signaling robustly specifies long-term repopulating HSCs among HSPCs along with coordination of downstream developmental differentiation choices^41^. Indeed, Fli-1^ROSAΔ^ HSPCs co-cultured with a vascular niche, displayed reduced Notch1 protein levels, and diminished activation of downstream Notch reporting signal (Fig. 3c, d). Hence, we propose that Fli-1 presets an HSPC activation program by producing angiokines such as Vegfa priming and inducing an adaptive and supportive Notch ligand expressing niche. Simultaneously, Fli-1 is transcriptionally coordinating an extracellular receptive sensory machinery for Notch signaling with the intracellular cooperative elements to propagate niche endorsed regenerative Notch activation signals for HSPCs (Fig. 3e). Further inspection of Notch1 genomic locus combining our multiome analyses indicated that Notch1 locus region contained regulatory elements which displayed Spi1 as well as Fli-1 binding activity. In Fli-1^ROSAΔ^ HSPCs these Spi1-targeted regions displayed reduced ATAC signal, representing reduced chromatin accessibility, yet presented an enhanced “footprinting” signal, indicating higher occupancy by a TF (Fig. 3f). This signifies that Spi1 may transcriptionally suppress Notch1 in order to prevent HSPC activation.

**Fig. 3:**
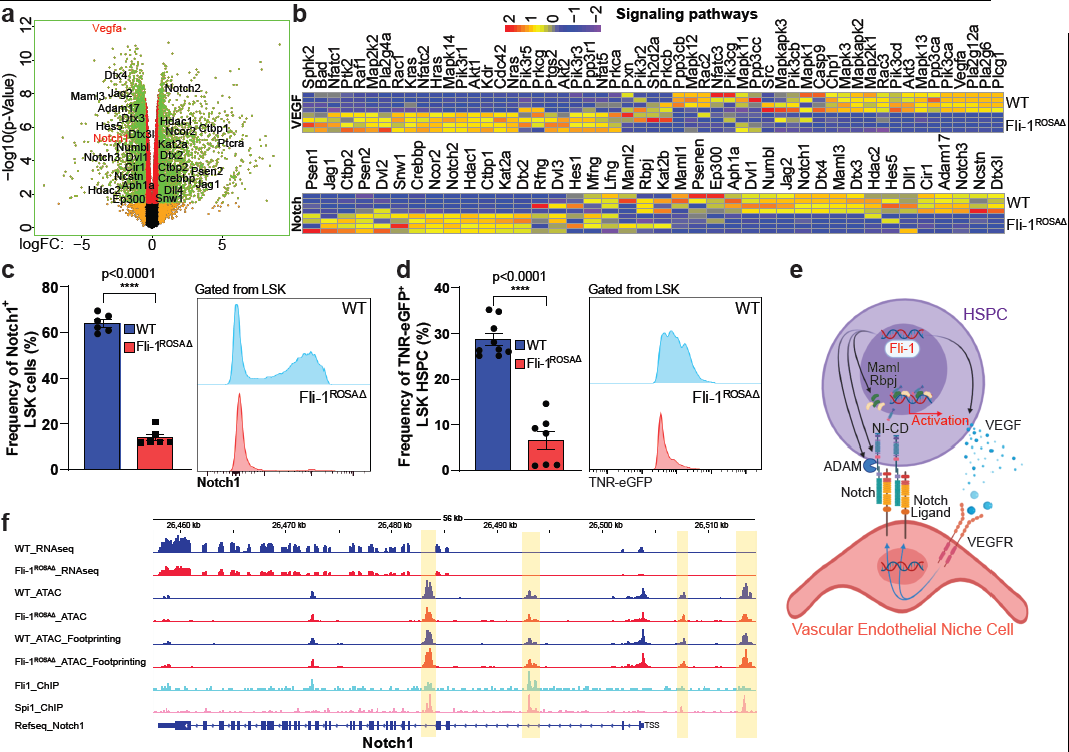
Fli-1 simultaneously poises regenerative activation programs via VEGF-A/Notch pathways crosstalk. **a,** Volcano plot of differentially expressed genes in Fli-1^ROSAΔ^ vs. WT HSPCs. Thresholds were set at |LogFC|>0.5 for x-axis and at FDR<0.05 for y-axis. Green and red color labeling is for genes with FDR<0.05 and green and yellow color labeling is for genes with |LogFC|>0.5. Genes belonging to the Notch signaling pathway and Vegfa are annotated to their localization on the heatmap. See also Table S1. **b,** Heatmaps for KEGG database extracted Notch/VEGF-A pathways elements across RNA-seq replicates of Fli-1^ROSAΔ^ and WT HSPCs. Scale bar and coloring represent the Z-score scaled by gene for Log counts per 10^6^ normalized by library size (trimmed mean of M-values). **c,** WT and Fli-1^ROSAΔ^ BM HSPCs were isolated and expanded. Harvested HSPCs were stained and analyzed by flow cytometry to determine frequency of Notch1 expressing LSK HSPCs. Representative Notch1 histogram plots gated from LSK HSPCs. Unpaired two tailed t-test was used; n=6 tissue donor mice per genotype, each point represents an average of 3-4 technical replicates. **d,** WT and Fli-1^ROSAΔ^ were bred with transgenic Notch reporter eGFP (TNR-eGFP) mice, to evaluate changes in downstream Notch signaling. BM LSK HSPCs were isolated and sorted from TNR-eGFP/ WT or Fli-1^ROSAΔ^ mice and expanded in co-cultures with a vascular niche layer. Frequency of LSK HSPCs positive for Notch signaling (eGFP^+^) as determined by flow cytometry at the end point of co-cultures. Representative flow histograms display TNR-eGFP signaling in WT (upper, blue) and Fli-1^ROSAΔ^ (lower, red) LSK HSPCs. Unpaired two tailed t-test was used; n=9 (WT) and n=7 (Fli-1^ROSAΔ^). **e,** Proposed model for Fli-1 endorsed HSPC activation via Notch/VEGF-A pathways mediated crosstalk with the vascular niche. Fli-1 in HSPCs, transcriptionally presets Notch pathway promoting external and internal elements while also transcriptionally promoting Vegfa expression. Vegfa is secreted to enlist an activation supportive vascular niche by stimulating ECs to elevate expression of Notch ligands. Reciprocal niche-mediated activation of Notch receptors presented by HSPCs, endorse downstream Notch signaling, promoting HSPC regenerative activation and expansion. **f,** IGV peaks plots for the genomic loci of Notch1 displaying peaks calling analyses for WT and Fli-1^ROSAΔ^ RNA-seq, ATAC-seq, ATAC-seq-“Footprinting”, Fli1-ChIP-seq, and Spi1-ChIP-seq. Yellow masking highlights regions displaying ATAC, “footprinting”, and Fli-1 + Spi1 TF binding activities. Note opposing correlation between ATAC to “footprinting” activities, among WT and Fli-1^ROSAΔ^ samples, for these regions, preferentially for the ones with stronger Spi1 binding activity.

Thus, we propose that Fli-1 transcriptionally orchestrates the sensing and adaptability to niche-derived Notch signals required to augment a regenerative activation program in HSPCs. Spi-1’s targeting of Notch1 may indicate for a possible transcriptional suppression of this pathway to preserve HSCs in a quiescent state and prevent the sensing of activation guiding signals arriving from the surrounding microenvironment.

## Induction of Notch1 signaling rescues expansion capacity of Fli-1 deficient HSPC

To determine whether HSPCs are receptive to a customized regenerative vascular niche, rather than purely acting autonomously, we bred Fli-1^ROSAΔ^ mice with mice carrying a conditional Cre inducible transgene of the intracellular component of the murine Notch1 gene lacking the c-terminal domain (N1-IC^iOE^). In this combined mouse line, upon tamoxifen induction there is a simultaneous deletion of Fli-1 alleles and induction of constitutively enforced Notch1 signaling activity (Fli-1^ROSAΔ^N1-IC^iOE^), mimicking the activation of Notch1 signaling by a stimulated niche. Early studies showed that Notch1 overexpression in HSPCs increases self-renewal while hindering differentiation rate, also skewing it towards a lymphoid output^39, 40^. Accordingly, Fli-1^ROSAΔ^N1-IC^iOE^ HSPCs demonstrated a rescue of HSPC expansion in co-culture with a vascular niche platform, preferentially expanding immature HSPCs, while still lacking megakaryocytic development (Fig. 4a, b **and** Extended Data Fig. 6a-c). Furthermore, transplantation of expanded Fli-1^ROSAΔ^N1-IC^iOE^ HSPCs revealed an engraftment rescue as these HSPCs contributed both to short-and long-term hematopoiesis, unlike Fli-1^ROSAΔ^ HSPCs which completely fail to repopulate recipient mice. Notably, transplanted Fli-1^ROSAΔ^N1-IC^iOE^ HSPCs exhibited an opposing differentiation pattern to WT HSPCs, during repopulation and recovery, reconstituting initially mostly lymphoid B-cell and T-cell output after short term engraftment but eventually exhibiting also restoration of the myeloid lineage, following long-term engraftment period (Fig. 4c **and** Extended Data Fig. 6d-f).

**Fig. 4:**
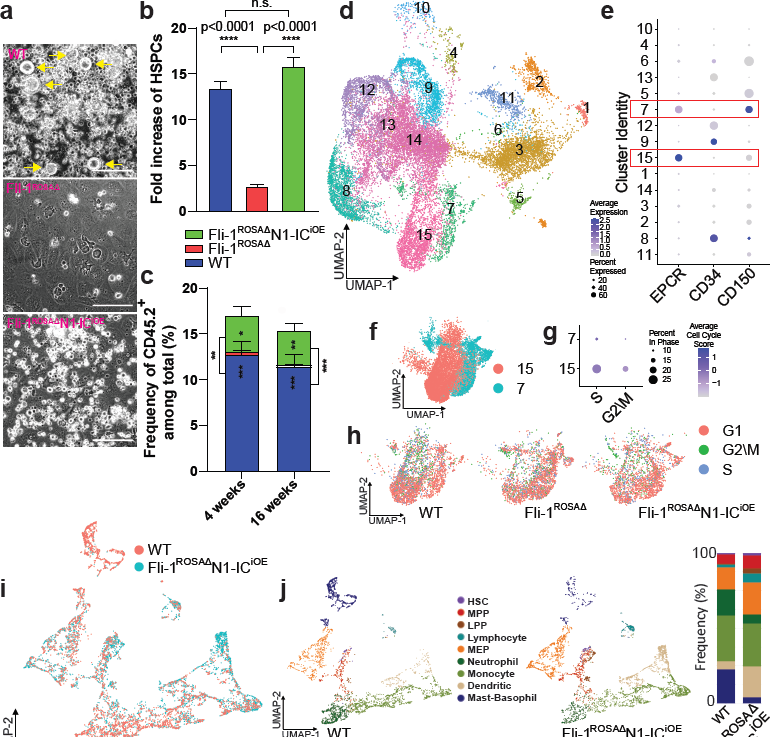
Enforcing Notch1 activation in Fli-1 deficient HSPCs restores regenerative expansion capacities and stem cell functionality. WT, Fli-1^ROSAΔ^, and Fli-1^ROSAΔ^ with conditionally inducible Notch1 internal component overexpression transgene (Fli-1^ROSAΔ^N1-IC^iOE^) BM LSK HSPCs were isolated and expanded as described in Fig. S5A. Harvested cells were analyzed by flow cytometry and/or flow sorted for LSK HSPCs which were competitively co-transplanted with congenic SJL BM cells into lethally irradiated congenic SJL recipient mice. **a,** Representative images of co-cultures at the end point before harvest. Yellow arrows indicate megakaryocytes. Note expansion of round hematopoietic cells in Fli-1^ROSAΔ^N1-IC^iOE^ co-cultures without any appearance of megakaryocytes. Bar = 100 µM. **b,** Frequency of LSK HSPCs was determined by flow cytometry and fold expansion was calculated. One-way ANOVA multiple comparisons was used; n=4 BM donor mice per genotype, with 2 technical replicates per donor. **c,** Frequency of chimerism indicating engraftment levels as determined by flow cytometry. One-way ANOVA multiple comparisons was used; n=8 recipient mice per genotype. **d-g,** WT, Fli-1^ROSAΔ^, and Fli-1^ROSAΔ^ with conditionally inducible Notch1 internal component overexpression transgene (Fli-1^ROSAΔ^N1-IC^iOE^) BM LSK HSPCs were isolated and expanded. After 48 hours of expansion in co-culture, cells were harvested and hematopoietic LSK HSPCs were sorted and applied for single cell RNA-seq analysis; n=4 per genotype “pooled” together. **d,** Dimensionality reduction by UMAP of single cell transcriptomes from co-cultured and sorted LSK HSPCs. **e.** Dot plot for the E-SLAM HSC markers EPCR, CD34, and CD150 per cluster identity. Dot plot size indicates percent expression in cluster and color intensity indicates average expression score. **f.** Distribution of clusters 7 and 15 in UMAP for HSC identified populations among LSK HSPC clusters based on Garnette and the Dot plot from panel (e). **g,** Dot plot for cell cycle scores (S phase and G2/M phases) per cluster identity. Dot plot size indicates percent expression in cluster and color intensity indicates average expression score. **h,** Cell cycle status classification in UMAP for clusters 7 and 15: WT (left UMAP), Fli-1^ROSAΔ^ (middle UMAP) and Fli1^ROSAΔ^N1-IC^iOE^ (right UMAP). **i-l,** BM cells were harvested, 16 weeks post engraftment of WT and “rescued” Fli-1^ROSAΔ^N1-IC^iOE^ long term repopulating HSCs and pooled from n=4 recipient mice (from 4 different donors) per genotype. CD45.2^+^/Lin^−^ cells were flow sorted and applied for single cell RNA-seq analysis. (HSC: hematopoietic stem cells, MPP: multi-potent progenitors, LPP: lymphoid-primed progenitors, MEP: megakaryocyte/erythroid progenitors). **i,** Dimensionality reduction by UMAP of single cell transcriptomes from donor-derived lineage-negative BM samples (CD45.2^+^/Lin^−^) following transplantation of WT or Fli1^ROSAΔ^N1-IC^iOE^ LSK HSPC. **j,** Cell type classification of hematopoietic sub-populations using Garnette: WT (left panel) and Fli1^ROSAΔ^N1-IC^iOE^ (right panel) samples in UMAP, and distribution of cell types between samples (bar plot). **k,** Distribution of WT and Fli1^ROSAΔ^N1-IC^iOE^ sample cells in UMAP among populations classified as multi-lineage stem/progenitor cell types (HSC, MPP, or LPP) using Garnette, and distribution of cell types between samples (bar plot). See also Table S5. **l**, Heatmaps of gene-set scores for the subset of multi-lineage stem and progenitor cell types (HSC, MPP, LPP) in UMAP (left panels), and violin plots of gene-set scores between samples (WT vs Fli1^ROSAΔ^N1-IC^iOE^) (right panels), for HSC molecular overlap signature genes (from Wilson *et al*. 2015) and engrafting long term HSC signature genes (from Rodriguez-Fraiticelli *et al*. 2020). Gene-set scores in violin plots are shown only for the subset of cells classified as HSC cell type. Wilcoxon Rank Sum Test was used to determine p-Values.

To study the dynamics of HSCs in co-culture *in vitro* conditions, we have sorted and single cell RNA-sequenced (scRNA-seq) expanding LSK HSPCs 48 hours post induction of Fli-1 deficiency. Following classification of heterogenous HSPC sub-clusters with combination of E-SLAM markers (EPCR^+^\CD34^−^\CD150^+^), we identified two HSC sub-clusters (#7 and 15#) (Fig. 4d, e). Among these two HSC sub-clusters, Fli-1 deficiency decreased the number of cells in cluster #15, reducing its cellular portion. Induction of Notch1 signaling in Fli-1 deficient HSPCs managed to effectively restore the subset of cells in cluster #15 (Extended Data Fig. 7a, b). Interpreting the genomic differences for these two clusters by GSEA for biological pathways, revealed very high and significant expression of genes involved in lysosome processing, which is essential for active maintenance of quiescent HSCs^42^, in cluster #7. Cluster #15 exhibited high expression of activation genes defined as stem cell pluripotency pathways and additionally higher expression of genes involved in activity of multiple signaling pathways such as Notch and WNT (Extended Data Fig. 7c). Indeed, cell cycle scoring of HSC sub-clusters revealed an activated state of cluster #15 and a quiescent state for cluster #7. Cycling cells disappear from cluster #15 following Fli-1 deletion and are restored upon enforced induction of Notch1 signaling (Fig. 4f-h **and** Extended Data Fig. 7a, b).

## Maintenance of Notch1 signaling in the absence of Fli-1 rescues HSC functionality

To validate the rescue of functional HSCs by niche restored signals, we isolated WT and Fli-1^ROSAΔ^N1-IC^iOE^ donor-derived immature CD45^+^/Lineage^negative^ BM cells at 4 months post-transplant and performed scRNA-seq analysis. Fli-1^ROSAΔ^N1-IC^iOE^ HSPCs displayed multipotent contribution to the more mature transcriptionally defined hematopoietic cell types in the lineage-negative marrow compartment, though with a relative skewing toward lymphoid cells and differential distribution among various myeloid populations compared with WT HSPCs (Fig. 4i, j). “Zooming in” on the portion of UMAP space representing transcriptionally defined HSCs, multi-potent progenitors (MPP), and lymphoid-primed progenitors (LPP), we observed that Fli-1^ROSAΔ^N1-IC^iOE^ HSPCs contribute to the HSC population at a similar frequency as WT HSPCs, with relative skewing towards the LPPs at the expense of the MPPs (Fig. 4k). Moreover, using previously published gene signatures derived from scRNA-seq data for functional HSCs^43^ with LTR engraftment potential^11^, we show that Fli-1^ROSAΔ^N1-IC^iOE^ HSPCs contain transcriptionally-defined functionally long-term engrafting HSCs (Fig. 4l), exhibiting some differential transcriptional output (**Supplementary Table 7**).

Thus, Fli-1 levied HSC regenerative activation program can be rescued by enforced expression of niche-propagated Notch signaling, confirming that niche-derived signals are essential to augment and support hardwired intrinsic stem cell programs.

## An active human HSPC signature associated with FLI-1 transcriptional activity categorizes cord blood HSCs versus quiescent mobilized HSCs

To translate the potential of Fli-1 dependent triggering of the stem cell activation programs, we focused our attention to adult human HSCs. One of the unmet needs and challenges in the field of human HSPC expansion for immune regenerative purposes is the capability to robustly expand the scarce population of human mobilized peripheral blood (mPB) HSPCs. Specifically in cases such as donor mobilization failure, donors with exhaustive BM, or to benefit HSC final yield prior/following genetic therapeutic manipulation^44^. In contrast, there is a significant advancement in the translational ability to expand human cord blood (CB) derived HSPCs^45–47^. To determine why mPB HSPCs are refractory to undergo *ex vivo* expansion, we performed scRNA-seq analysis of sorted CD45^+^CD34^+^ HSPCs, from both CB and mPB origins, following two days in co-culture with a vascular niche platform to prime the regenerative expansion of these cells. Higher diversity and HSPC heterogeneity were observed amongst CB CD34^+^ cells, which also exhibited an actively cycling state in comparison to mPB CD34^+^ cells that remained unresponsive to our regenerative expansion niche system (Fig. 5a-d). GO reactome analysis of differentially expressed genes in the HSC containing cluster between CB and mPB, exhibited activation of mitotic cycling pathways amongst CB cells. By contrast, mPB cells displayed inflammatory related pathways, mainly of quiescence associated interferon signaling (Extended Data Fig. 8a **and Supplementary Table 8**).

**Fig. 5:**
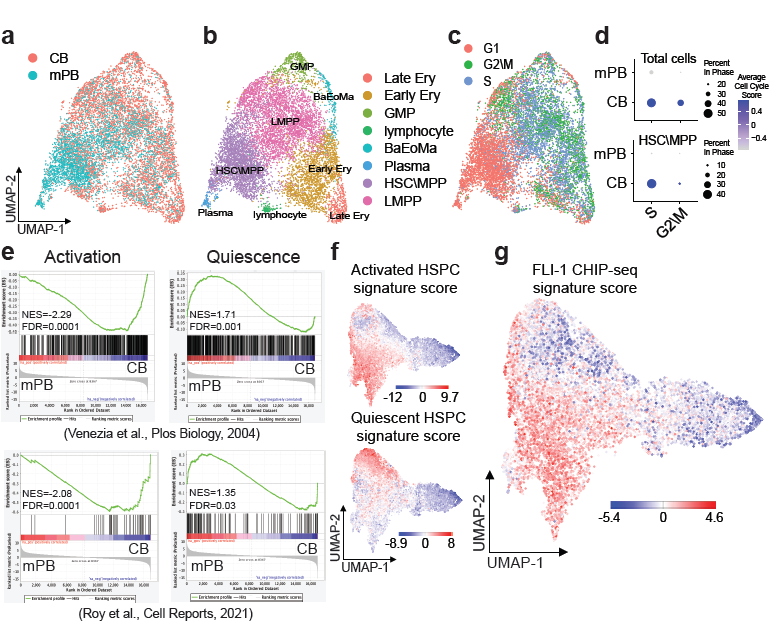
Transcriptional comparison distinguishes distinct activation/quiescence states for neonatal and adult HSPCs. **a-d,** Sorted CD45^+^/CD34^+^ HSPC from cord blood (CB) and adult mobilized peripheral blood (mPB) sources, were co-cultured on top of a vascular niche for 48h to encourage HSPC activation. Next, cells were harvested and CD45^+^/CD34^+^ HSPC were sorted and applied for single cell RNA-seq analysis. **a-c,** Dimensionality reduction by UMAP of single cell transcriptomes from co-cultured and sorted CD45^+^/CD34^+^ HSPC, displaying cell distribution by CB and mPB source (a), cluster identity (b), and single cell cycle phase (c). **d,** Dot plot for cell cycle scores (S phase and G2/M phases) per cellular source (CB or mPB), for total cells in analysis (upper panel) and for HSC\MPP cluster (lower panel). Dot plot size indicates percent expression in cluster and color intensity indicates average expression score. **e,** To highly enrich for HSCs, the mononuclear fraction from CB (n=3) and mPB (n=4) sources was isolated, labeled, and sorted for CD45^+^/CD34^+^/CD38^−^/CD45RA^−^/CD90^+^/CD49f^+^. Sorted cells were lysed and processed for RNAseq analysis. GSEA analysis plots for HSC activation signatures (left panels) and for HSC quiescence signatures (right panels) acquired from Venezia et al., 2004 (upper panels) and from Roy et al., 2021 (lower panels), showing a positive enrichment for the activation signatures in CB HSCs and positive enrichment for quiescence signatures in mPB HSCs. **f, g,** UMAP projections of the single cell ATAC-seq enrichment analysis for sorted HSPCs of all 10 signatures identified in Takayama et al., 2021. Colors indicate the degree of enrichment in each cell (blue, depleted; red, enriched). Scale bar indicates enrichment Z-score. **f,** Enrichment for activated HSPCs (upper panel) and quiescent HSPCs (lower panel) signatures defined in in Takayama et al., 2021., as calculated by chromVAR. **g,** Enrichment for called peaks from the FLI-1 HSPC ChIP-seq (Fli-1 signature) data set (Beck et al., 2013) overlayed on single cell ATAC-seq HSPC data set (Takayama et al., 2021), as analyzed by chromVAR.

To unravel if these aberrant HSPC responses were primed by the signals emanating from the niche or were a pre-existing cell intrinsic program, we have analyzed RNA-seq of sorted and highly purified HSCs from CB and mPB origins. Repeatedly, CB HSCs revealed a hallmark genomic signature associated with activation of multiple metabolic pathways and myc targets, indicating for their cycling potential. By contrast, the mPB HSCs displayed mainly inflammatory and interferon signaling pathways along with activation of TGFβ signaling and Trp53 pathways (Extended Data Fig. 8b **and Supplementary Table 9**). Further applying previously defined gene lists^48, 49^ for HSC activation and quiescence states, we confirmed CB HSC active versus mPB HSC quiescent natures (Fig. 5e). Next, to ascertain human HSCs FLI-1’s role as activation factor, we reanalyzed previously described scATAC-seq data set for human HSPCs^50^ together with reported FLI-1’s ChIP-seq peaks signature in human HSPCs^51^. After assigning the active HSC and quiescent HSC signature scores as defined by Takayama et al^50^, we applied the FLI-1 ChIP-seq signature score to define which HSCs exhibited accessible regions targeted by FLI-1 as indication for FLI-1 activity. Almost a complete overlap was observed between the signatures for FLI-1 ChIP and active HSCs (Fig. 5f, g), confirming the association between FLI-1 transcriptional activity to human HSC activation as well. Thus, enforcing FLI-1 mediated chromatin remodeling may have the potential to override adult mPB HSC quiescence barrier and promote the adaptation to a regenerative niche platform.

## Transient FLI-1 overexpression activates mobilized adult human HSPCs

Modified mRNA is an emerging non-immunogenic, efficient, and transient tool for safe induction of protein expression^52^, with high relevance for potential oncogenes that can direct cell activation, such as in the case of Pkm2-induced cardiac regeneration^53^. We have applied modified-mRNA for FLI-1 transient expression in sorted human CD34^+^ HSPCs, followed by a co-culture with a regenerative vascular niche platform to drive HSPC expansion prior to a pre-clinical study of transplantation into immunodeficient mice (Fig. 6a). As expected, co-culturing mPB HSPCs with a vascular niche exhibited poor expansion potential in comparison to CB HSPCs (Fig. 6b, c). Following transduction of FLI-1 modified-mRNA no beneficial effect was observed for CB HSPC expansion probably due to their already active nature, except for the more primitive CD34^+^CD38^neg^ HSPC, that preserve a latent activation resistant sub-population of HSCs^54^ (Fig. 6b). Contrary, mPB HSPCs displayed a significant expansion benefit across mature hematopoietic and immature HSPC sub-populations, following FLI-1 modified mRNA transduction, achieving augmented expansion levels comparable to unstimulated CB HSPCs (Fig. 6b-d). Further transplantation studies of expanded control and FLI-1 transduced mPB HSPCs, demonstrated for the later a superior engraftment and human hematopoietic reconstitution capacity in the peripheral blood (PB), the spleen, and the BM of immunodeficient mice (Fig. 6e), with higher frequency of human HSPCs present in the murine BM (Fig. 6f, g). Secondary transplantation of BM cells confirmed the higher presence of human HSCs in the BM of primary recipient immunodeficient mice engrafted with FLI-1 transduced mPB HSPCs (Fig. 6h).

**Fig. 6:**
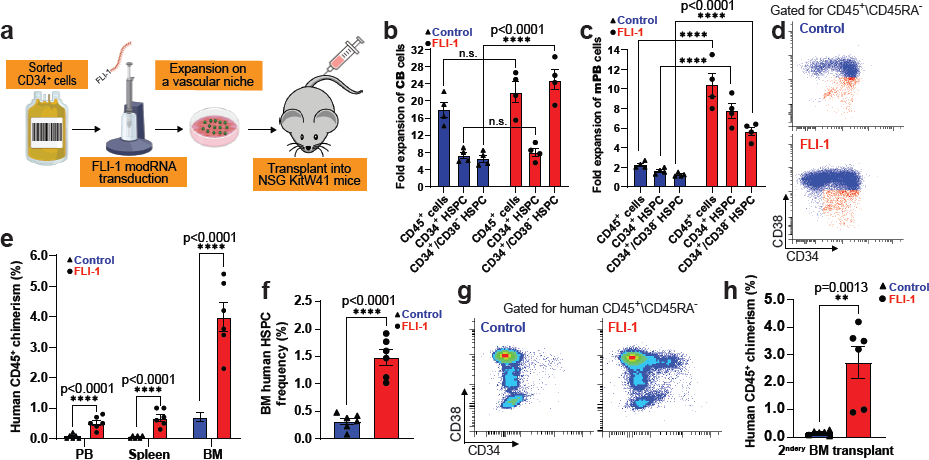
Activation of adult human mobilized HSPC via transient overexpression of FLI-1. **a,** Schema of experimental design. Sorted human CD34^+^ HSPCs from cord blood (CB) or mobilized peripheral blood (mPB) sources were transduced by electroporation with FLI-1 modified-RNA molecules (2 µg FLI-1 modRNA per 10^5^ cells) and expanded for 1 week on top of E4orf1 vascular niche cells in a sub-optimal ratio of 1:3 (HSPCs:ECs). Next, expansion co-cultures were analyzed and transplanted into immunodeficient NSG KitW41 mice without myeloablative preconditioning. **b**, Fold expansion of CB derived hematopoietic subtypes after 1 week in co-culture following transduction with FLI-1 (red) or control (blue) modified-RNA. Unpaired two tailed t-test was used; n = 4 CB donors. Each mark represents the averaged triplicate (n = 3 technical repeats) per donor. **c**, Fold expansion of mPB derived hematopoietic subtypes after 1 week in co-culture following transduction with FLI-1 (red) or control (blue) modified-RNA. Unpaired two tailed t-test was used; n = 4 mPB donors. Each mark represents the averaged triplicate (n = 3 technical repeats) per donor. **d,** Representative flow dot plots of mPB HSPC analysis post 1 week of expansion in co-culture following transduction with FLI-1 or control modified-RNA. The population of CD34^+^\CD38^neg^ HSPCs is colored in red. **e,** Human CD45 chimerism analysis in peripheral blood (PB), spleen, and bone marrow (BM) of NSG KitW41 mice. Tissues were harvested and analyzed by flow cytometry 16 weeks post transplantation. Unpaired two tailed t-test was used; n = 6 mPB donors. **f,** Frequency of BM engrafted human HSPCs as determined by flow cytometry 16 weeks post transplantation. Unpaired two tailed t-test was used; n = 6 mPB donors. **g,** Representative flow dot plots for BM engrafted human CD34^+^ and CD34^+^\CD38^neg^ HSPCs acquired 16 weeks post transplantation. Unpaired two tailed t-test was used; n = 6 mPB donors. **h,** Total BM from primary NSG KitW41 recipient mice was transplanted into secondary NSG KitW41 recipient mice without myeloablative preconditioning. Human CD45 chimerism in the BM of recipient mice was determined by flow cytometry 16 weeks post secondary transplantation. Unpaired two tailed t-test was used; n = 6 mPB donors.

Hence, transient overexpression of activation factors such as FLI-1 can be applied to direct the expansion of quiescent mPB HSCs, that ignore expansion signals delivered from a regenerative vascular niche platform (Extended Data Fig. 9). This approach most probably can be leveraged to promote mPB HSC expansion with other existing platforms, currently effective only for CB HSC.

## Discussion

Herein, we have probed the two-way relationship between the stem cell and its vascular niche, by unraveling a new paradigm in the physiology of adult hematopoiesis. We demonstrate that the transition from quiescence to active regenerative hematopoiesis is dependent on Fli-1 enabling HSCs to sense, relay, adapt, and to execute a niche triggered activation program. Fli-1 empowers HSPCs with the capacity to interpret the pertinent niche signals destined for the transition into a hyperactive state. Vascular ECs, representing prototypical niche cells, respond to Fli-1-governed VEGF-A counter signal from HSPCs to customize a Notch-driven niche response towards support of regenerative hematopoiesis. Meanwhile, other niche cells such as osteoprogenitors, respond to cholinergic regenerative signals, to customize niche derived cues that preserve the quiescent HSC pool^55^. Thus, Fli-1 intrinsic activity in HSPCs enables the crosstalk with expansion designated niche cells, allowing us to “molecularly eavesdrop” on the reciprocal activation signals sustaining hematopoietic active regeneration and post-insult immune recovery.

Harnessing our findings, we were able to activate human quiescent mPB HSPCs with FLI-1 modified-RNA, and to boost their expansion on a vascular niche platform to the matching level of naturally activated human CB HSPCs. Employing modified-RNA delivery into HSPCs can be benchmarked as a clinically safe technology allowing transient protein upregulation by cells, without provoking any potential pro-leukemic transformations. Enhanced engraftment levels of FLI-1 stimulated human cultures presents a therapeutic solution for patients with insufficient numbers of transplantable mPB HSPCs. The revelations gathered from our study, in which synthetic modified-RNA technology enable transient overexpression and activation of a factor or a combination of factors, can be harnessed to steer quiescent HSCs for optimal *ex vivo* expansion in a pre-transplantation clinical setting, as in the cases of poorly-mobilizing donors suffering from exhausted chemotherapy/irradiated stressed BM, or co-morbidities, including diabetes^44^. Mostly, it could benefit and significantly improve therapeutical efforts of autologous based HSC gene therapy, as in the case of β-thalassemia, sickle cell, and other rare diseases, by tackling currently limiting attributes^56^. HSC release from quiescence may enhance the transduction rate of the engineered vector, and may further mitigate p53-mediated clonal shrinking of gene edited HSCs^57^, and to expand the pool of successfully transduced HSCs, increasing the final yield of patient-engrafted engineered HSCs.

Finally, our findings set forth the transformative notion of a co-adaptive and responsive rather than a dictating niche role. The notion of molecular hubs transcriptionally presetting cellular genetic programs, masterminding distinct immune stem and progenitor cell states, may apply globally to varieties of normal and malignant stem/progenitor and niche cell types. This may further reveal a myriad of specialized niche types designated for diverse stem/progenitor cell intrinsic transcriptional programs.

## Acknowledgment

We are grateful to the American Society of Hematology (ASH)-European Hematology Association (EHA) Translational Research Training in Hematology (TRTH) joint program for the valuable support and guidance of Dr. Tomer Itkin and this study. Drs. Ying Liu and Jesus M. Gomez Salinero are New York Stem Cell Foundation-Druckenmiller Fellows. Dr. Yang Lin is NYSTEM fellow. The authors would like to thank members of Weill Cornell Medicine’s Genomic Core facility for all their tremendous help and assistance. Illustrations in this manuscript were partially or entirely created with BioRender.com.

## Author Contribution

TI and SR conceived and designed the study, supervised experiments and analysis, performed experiments, analyzed data and wrote the manuscript. YL, CRB, YL, JMGS, FG, JHS performed experiments and assisted in data analysis. BH designed and together TI analyzed scRNA-seq data. JED, SZX, VV, AM, KBK designed, performed, and analyzed bulk RNA-seq experiments of human CB and mPB HSPCs. JAS designed and together with NS performed intravital imaging experiments and analyzed data. SH designed and wrote scripts, and with TI performed bioinformatic analyses of bulk RNA-seq, ATAC-seq, multiple types of ChIP-seq, scRNA-seq, and single nuclei multiome data. SZJ assisted in design and analysis of ATAC-seq related experiments. DR supervised and mentored SH. CT, GB, SH, TI designed performed and analyzed multiome snRNA/ATAC-seq analysis. RS designed and performed microscopy experiments and analyzed imaging data. JZX assisted with design and performance of all sequencing analysis. LZ designed and synthesized modified-mRNA and assisted in design of modified-mRNA related experiments.

## Data and Materials Availability

All codes and multi-omics data will be provided upon request from the corresponding authors.

## Address correspondence to

Tomer Itkin, Department of Medicine, Division of Regenerative Medicine, Ansary Stem Cell Institute, 413 East 69th Street, Room A-869, New York, New York 10021, USA. Phone: 212.746.2017.

Shahin Rafii, Department of Medicine, Division of Regenerative Medicine, Ansary Stem Cell Institute, 413 East 69th Street, Room A-863A, New York, New York 10021, USA. Phone: 212.746.2070;

## Supplementary Materials

### Methods

#### Supplementary Tables 1-10

**Supplementary Table 1:** snRNA-seq differential gene expression in distinct HSPC sub-clusters.

**Supplementary Table 2:** Full heatmap representation of differential ChromVAR motif activity in distinct HSPC sub-clusters by averaged Z-score, all predicted TF motifs names are presented.

**Supplementary Table 3:** snATAC-seq differential ChromVAR motif activity in distinct HSC/MPP sub-clusters and snRNA-seq differential gene expression in distinct HSC/MPP sub-clusters. GO for biological processes is presented for each set of snRNA/ATAC-seq analyses.

**Supplementary Table 4:** RNA-seq differential gene expression in Fli-1^ROSAΔ^ vs. WT HSPCs.

**Supplementary Table 5:** ATAC-seq differential gene peaks calling in Fli-1^ROSAΔ^ vs. WT HSPCs.

**Supplementary Table 6:** H3K27ac ChIP-seq differential gene peaks calling in Fli-1^ROSAΔ^ vs. WT HSPCs.

**Supplementary Table 7:** scRNA-seq differential gene expression in Fli-1^ROSAΔ^ vs. WT HSPC population.

**Supplementary Table 8:** scRNA-seq differential gene expression in CB HSC-MPP cluster vs. mPB HSC-MPP cluster.

**Supplementary Table 9:** RNA-seq differential gene expression in CB HSC svs. mPB HSC.

#### Supplementary Videos 1-5

**Supplementary Video 1: Intravital representative video of BM homed HSPCs.** DiD labeled Fli-1^ROSAΔ^ HSPC (red arrow) and autofluorescence (red). DiI labeled WT HSPC (white arrow), autofluorescence, and Dextran labeled blood vessels (blue). Bone - second harmonic generation of collagen (green). Bar = 50 µM. Z-step size = 3 µM.

**Supplementary Video 2: Intravital representative video of vessel wedged HSPCs.** DiI labeled Fli-1^ROSAΔ^ HSPCs (yellow arrows), autofluorescence, and Dextran labeled blood vessels (blue). Bone - second harmonic generation of collagen (green). Bar = 50 µM. Z-step size = 3 µM.

**Supplementary Video 3: Representative video of expending HSPCs in culture.** WT (left panel) and Fli-1^ROSAΔ^ (right panel) BM HSPCs were isolated and introduced into co-culture with E4orf1 vascular niche cells as described in Fig. S5A. Time tab indicates the hours that passed since the addition of 4-OHT into cultures. Bar = 500 µM.

**Supplementary Video 4: Representative video of expending HSPCs in culture.** WT (left panel) and Fli-1^ROSAΔ^ (right panel) BM HSPCs were isolated and introduced into co-culture with E4orf1 vascular niche cells as described in Fig. S5A. Time tab indicates the hours that passed since the addition of 4-OHT into cultures. Bar = 100 µM.

**Supplementary Video 5: Representative video of human adult mPB HSPCs in culture** Transduced Control (upper panel) and FLI-1 modified-RNA (lower panel) human adult mPB HSPCs were isolated and introduced into co-culture with E4orf1 vascular niche cells. Time tab indicates the hours that passed since introduction of HSPCs into co-cultures. Bar = 200 µM.

#### Extended Data Figures 1-9

**Extended Data Fig. 1:**
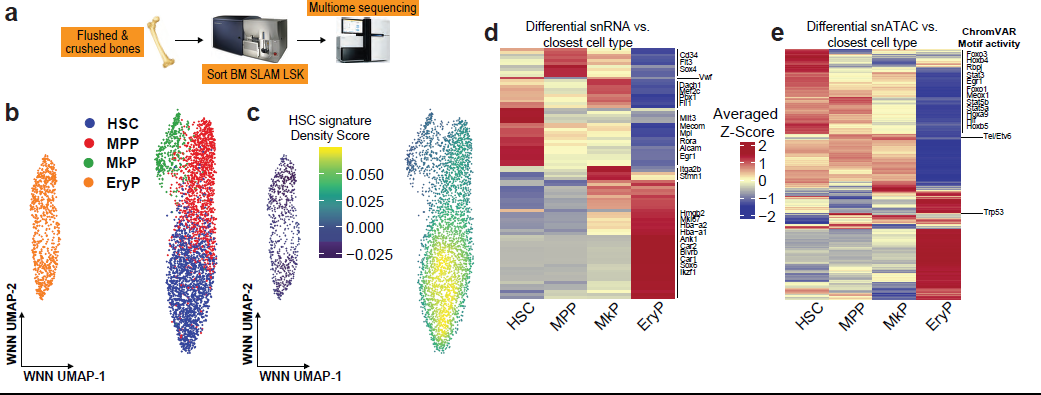
SLAM LSK HSPCs analysis classifies distinct HSPC clusters with unique RNA and motif activity signatures. **a,** Schema for analysis of BM primitive SLAM LSK HSPCs. Murine femurs and tibias were harvested, flushed, and crushed, to collect maximal yield of bone and marrow cells. BM labeled cells were flow sorted for the SLAM LSK markers: Live\Ter-119^neg^\Lineage^neg^\Sca-1^+^\c-Kit^+^\CD150^+^\CD48^neg^. Next, combined multiome single-nuclei RNA/ATAC (snRNA/ATAC) sequencing analysis was performed. **b,** Weighted nearest neighbor (WNN) UMAP with hematopoietic stem and progenitor cell type annotation for hematopoietic stem cell (HSC), multi-potent progenitor (MPP), megakaryocyte progenitor (MkP), and erythrocyte progenitor (EryP) sub-cluster representation. **c,** Engrafting LTR-HSC transcriptional signature (from Rodriguez-Fraiticelli *et al*. 2020) assigned on a WNN UMAP space. **d,** Heatmap representation of differential transcriptional nuclei output from distinct HSPC sub-clusters by averaged Z-score, with selected genes presented. **e,** Heatmap representation of differential ChromVAR motif activity in distinct HSPC sub-clusters by averaged Z-score, with selected TF motifs presented.

**Extended Data Fig. 2:**
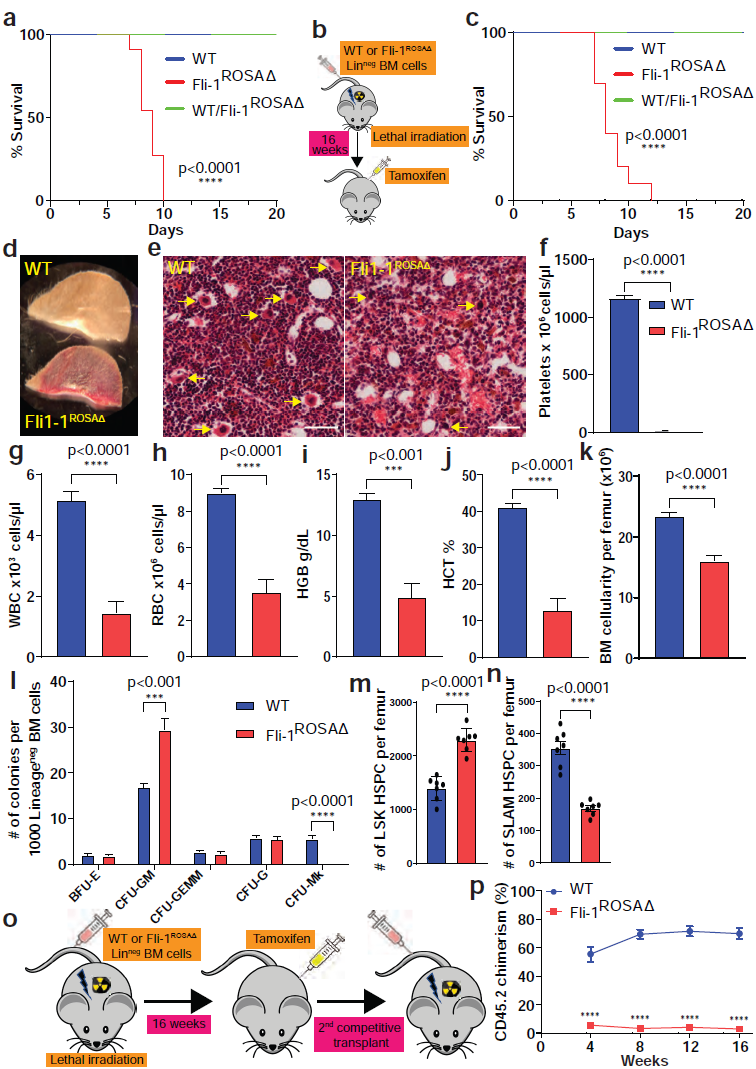
Hematopoietic Fli-1 deficiency leads to thrombocytopenia and to HSC regenerative failure. **a,** Survival plot for WT, Fli-1^ROSAΔ^, and WT/Fli-1^ROSAΔ^ (heterozygous) mice, following the beginning of tamoxifen induction (1^st^ injection on day 0). Gehan-Breslow-Wilcoxon test was used; n=9 (WT), n=11 (Fli-1^ROSAΔ^), and n=13 (WT/Fli-1^ROSAΔ^). **b,** Illustration of the working model to generate hematopoietic specific and inducible Fli-1^ROSAΔ^ mice. WT mice are lethally irradiated (475 rad + 475rad with 5 hours interval) and are transplanted with 2×10^5^ Lineage depleted (Lin^neg^) BM cells from either WT or Fli-1^ROSAΔ^ mice. Four months (16 weeks) post-transplant (full hematopoietic recovery) mice are administrated with tamoxifen to induce Fli-1 gene deletion. **c,** Survival plot for WT mice with either WT, Fli-1^ROSAΔ^, and WT/Fli-1^ROSAΔ^ (heterozygous) hematopoietic cells, following the beginning of tamoxifen induction (1^st^ injection on day 0). Gehan-Breslow-Wilcoxon test was used; n=9 (WT), n=10 (Fli-1^ROSAΔ^), and n=11 (WT/Fli-1^ROSAΔ^). **d,** Representative image of ear cuts from WT (upper) and Fli-1^ROSAΔ^ (lower) mice. **e,** A representative H&E image of femoral BM section from WT and Fli-1^ROSAΔ^ mice at day 7 post induction (n=5 mice per genotype). Bar = 50 µM. Yellow arrows indicate WT and abnormal Fli-1^ROSAΔ^ megakaryocytes. **f-j,** At day 7 post induction blood was collected and hematological parameters were scored using ADVIA 120. Platelet counts for WT and Fli-1^ROSAΔ^ mice are presented per 1 µL of blood. White blood cell (WBC) and red blood cell (RBC) counts are presented per 1 µL of blood. Hemoglobin (HGB) is presented in grams per deciliter. Hematocrit (HCT) frequency in blood is presented. Unpaired two tailed t-test was used; n=5 per WT or Fli-1^ROSAΔ^. **k,** At day 7 post induction femurs were flushed, stained with Turk’s solution and WBC were counted and determined using hemacytometer. Unpaired two tailed t-test was used; n=7 mice per genotype. **l,** Number of scored BM derived colonies per sub-type for WT and Fli-1^ROSAΔ^ mice at day 7 post induction. Multiple unpaired t-tests were performed; n=7 per WT or Fli-1^ROSAΔ^ with each n representing an average of two replicates. **m, n,** Number of BM LSK HSPC and SLAM LSK HSPC as determined by flow cytometry and normalized per WBC counts per femur at day 7 post induction. Unpaired two tailed t-test was used; n=7 per WT or Fli-1^ROSAΔ^. **o,** Illustration of the working flow to generate hematopoietic specific and inducible Fli-1^ROSAΔ^ mice, followed by Fli-1 knockout induction, and 2^ndery^ transplant. WT mice are lethally irradiated (475 rad + 475rad with 5 hours interval) and are transplanted with 2×10^5^ Lineage depleted (Lin^neg^) BM cells from either WT or Fli-1^ROSAΔ^ mice. Four months (16 weeks) post-transplant (full hematopoietic recovery) mice are administrated with tamoxifen to induce Fli-1 gene deletion. Seven days post induction bones were harvested, LSK HSPCs were sorted and competitively co-transplanted with congenic SJL BM cells into lethally irradiated congenic SJL mice. **p,** On day 7 post induction WT or Fli-1^ROSAΔ^ BM LSK HSPC were isolated by flow cytometry sorting (n=8 per genotype) and co-transplanted competitively with congenic SJL total BM cells into lethally irradiated congenic SJL recipient mice. Every 4 weeks peripheral blood was harvested and chimerism frequency was determined by flow cytometry. Multiple unpaired t-tests were performed. ****p<0.0001.

**Extended Data Fig. 3:**
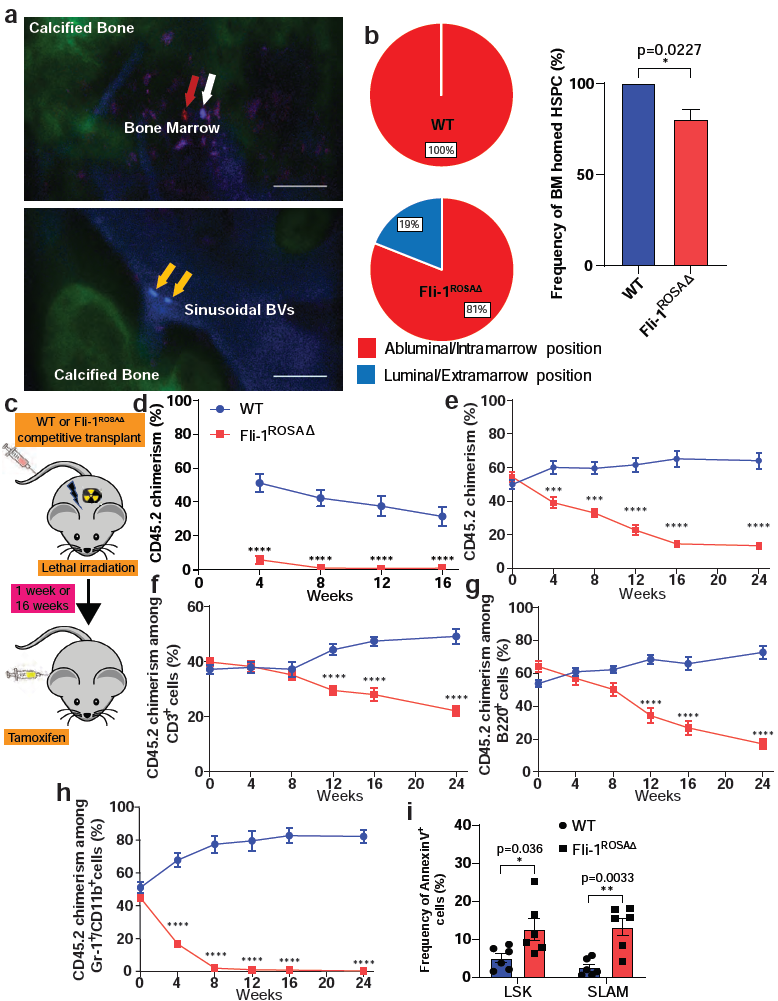
Fli-1 deficient HSPC regenerative failure is due to hampered hematopoiesis rather than due to homing defects. BM WT and Fli-1^ROSAΔ^ Lineage depleted cells were harvested and plated on top of a vascular niche layer. After 48 hours 4-hydroxytamoxifen (4-OHT) was administrated to induce Fli-1 deletion. After additional 48 hours LSK HSPCs were sorted and frozen for further experiments. Thawed WT and Fli-1^ROSAΔ^ HSPCs were labeled with either DiI or DiD dyes (10µM) (swapping dye labeling per genotype between experiments was performed to ensure no dye related effects). Recipient mice were sub-lethally irradiated (300 rad + 300 rad with 5 hours interval) 24 hours before transplantation. WT and Fli-1^ROSAΔ^ HSPCs were co-injected into recipient mice and 24 hours later mouse calvarias were imaged and cell luminal or abluminal location was determined using multiphoton/confocal laser scanning video-rate microscopy. **a,** Upper representative image – DiD labeled Fli-1^ROSAΔ^ HSPC (red arrow) and autofluorescence (red). DiI labeled WT HSPC (white arrow), autofluorescence, and Dextran labeled blood vessels (blue). Bone -second harmonic generation of collagen (green). Lower representative image – DiI labeled Fli-1^ROSAΔ^ HSPCs (yellow arrows), autofluorescence, and Dextran labeled blood vessels (blue). Bone - second harmonic generation of collagen (green). Note vessel wedged Fli-1^ROSAΔ^ HSPCs at lower panel image. Bars = 50 µM. See also Movies S1 and S2. **b,** Bars and pies presenting the portion of HSPCs located in abluminal/intramarrow (homed) or luminal/extramarrow (not homed) locations. Unpaired two tailed t-test was used; n=3 biological repeats of experiment per genotype with n=55 single HSPC events imaged and recorded. **c-i,** Lin^−^ WT or Fli-1^ROSAΔ^ BM cells mixed with congenic SJL BM cells (1:1 ratio) were co-transplanted into congenic SJL lethally irradiated recipient mice. **c,** Illustration of the working flow to generate mice with mixed chimeric hematopoietic system containing both WT and inducible Fli-1^ROSAΔ^ hematopoietic cells, to allow the study of Fli-1 deficient HSCs in a microenvironment containing WT hematopoietic and stromal cells. Lin^−^ WT or Fli-1^ROSAΔ^ BM cells mixed with congenic SJL BM cells (1:1 ratio) were co-transplanted into congenic SJL lethally irradiated recipient mice. Seven days post-transplant (to ensure successful hematopoietic BM lodgment), and four months post-transplant (to ensure complete hematopoietic BM recovery), tamoxifen was administrated for induction. **d,** Seven days post-transplant, to ensure successful hematopoietic BM lodgment, tamoxifen was administrated for induction. Every 4 weeks post induction peripheral blood was harvested and levels of chimerism were determined by flow cytometry. Multiple unpaired t-tests were performed; n=15 per WT and n=10 per Fli-1^ROSAΔ^. ****p<0.0001. **e-h,** Four months post-transplant, to ensure complete hematopoietic BM recovery, tamoxifen was administrated for induction. On day 0 Prior to induction and at indicated time points post induction peripheral blood was harvested and levels of chimerism were determined by flow cytometry among total hematopoietic cells, and myeloid/lymphoid lineage specific cells. Multiple unpaired t-tests were performed; n=8 per genotype. ***p<0.001, ****p<0.0001. **i,** Eight weeks post induction, BM cells were harvested and apoptosis frequency among WT or Fli-1^ROSAΔ^ LSK and SLAM LSK HSPCs was determined by flow cytometry. Unpaired two tailed t-test was used; n=6 per genotype.

**Extended Data Fig. 4:**
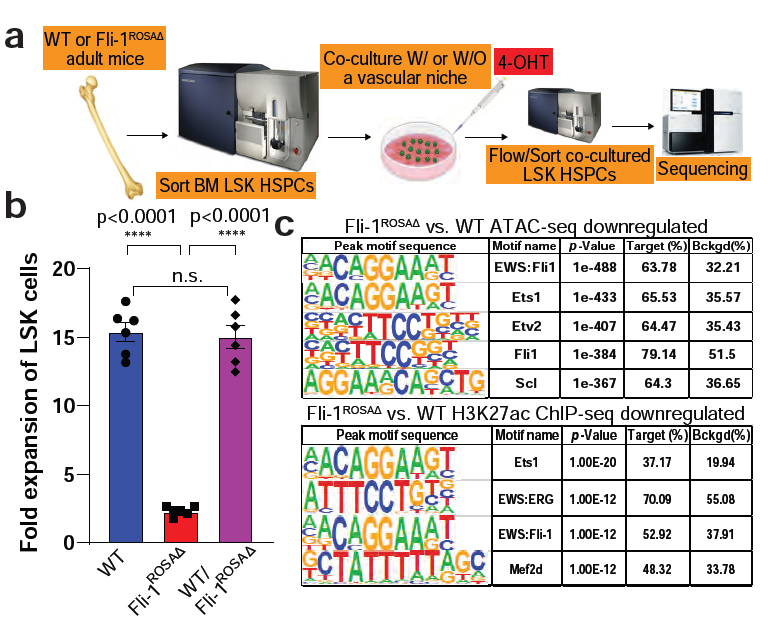
Fli-1^ROSAΔ^ HSPCs exhibit deactivation of regenerative programs. **a,** Illustrated working flow for studies of adult HSPC expansion in co-culture with a vascular niche. Bones are harvested from adult WT or Fli-1^ROSAΔ^ mice, flushed, crushed, and LSK HSPCs are isolated by flow sorting. Next, HSPCs are co-cultured with a layer of a vascular niche for 48h to allow recovery and induction of cellular expansion program. After 48h 4-OHT is added to the co-cultures to induce Fli-1 deletion, and co-cultures are kept for additional 6 days, supplemented with cytokine containing serum free media every other day. At the end point co-cultures are analyzed by flow cytometry and/or flow sorted for LSK HSPCs, which are further applied for different omics sequencing, and bioinformatical analyses. **b,** BM LSK HSPCs were sorted and introduced into a serum free co-culture condition with a vascular niche layer and with cytokine supplements. Frequency of LSK HSPCs was determined by flow cytometry and fold expansion was calculated. One-way ANOVA multiple comparisons was used; n=6 BM donor mice per genotype, each point represents an average of 3 technical replicates. **c,** Transcription factor binding site motif enrichment analysis for downregulated ATAC- and H3K27ac ChIP-seq peak sites in Fli-1^ROSAΔ^ vs. WT HSPCs. See also Tables S2 and S3.

**Extended Data Fig. 5:**
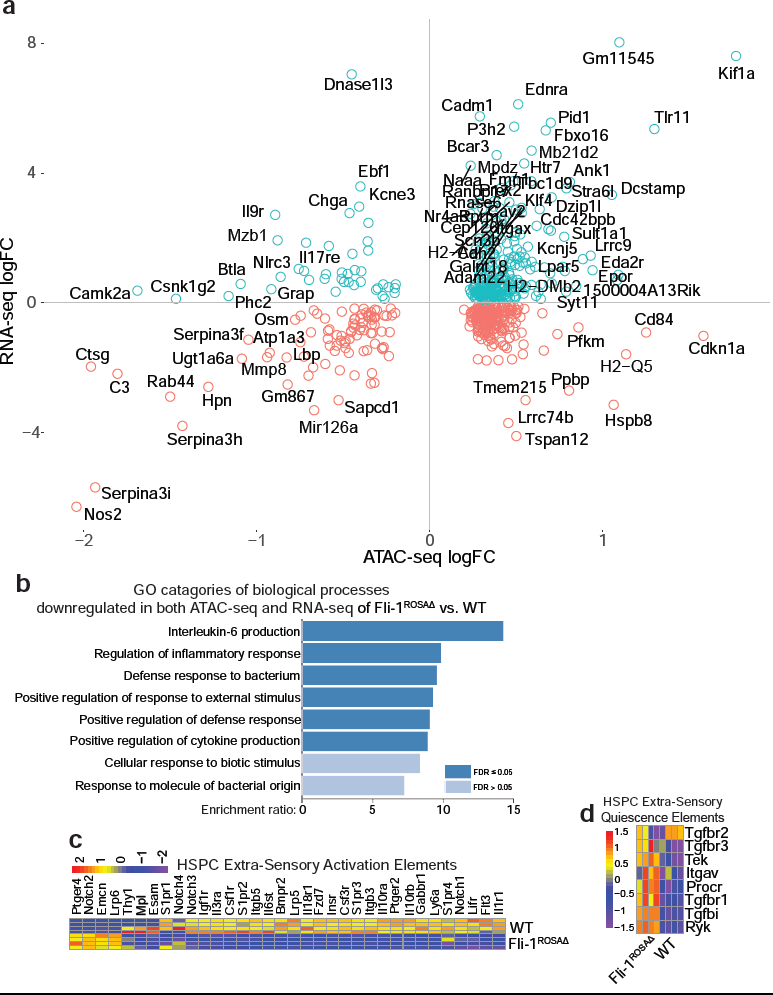
Intersecting transcriptional and chromatin accessibility data reveals HSPC sensory and cytokine production deficiencies in the absence of Fli-1. **a,** Intersection dot plot of RNA-seq and ATAC-seq in Fli-1^ROSAΔ^ vs. WT HSPCs. Each dot represents the transcriptional and chromatic accessibility statuses per gene. For gene labeling of few selected dots, thresholds FDR<0.05 for both RNA- and ATAC-seq, |LogFC|>2.5 for RNA-seq, |LogFC|>0.75 for ATAC-seq, and |Distance to TSS| < 500 parameters were set. **b,** Gene ontology (GO) categories of biological processes enrichment for significantly downregulated genes in both RNA- and ATAC-seq analyses of Fli-1^ROSAΔ^ vs. WT HSPCs (lower left quadrant in (a)). Significance by FDR value is indicated. **c, d,** Heatmaps for selected HSC extra-sensory activation (c) and quiescence (d) elements across RNA-seq replicates of Fli-1^ROSAΔ^ and WT HSPCs. Scale bar and coloring represent the Z-score scaled by gene for Log counts per 10^6^ normalized by library size (trimmed mean of M-values).

**Extended Data Fig. 6:**
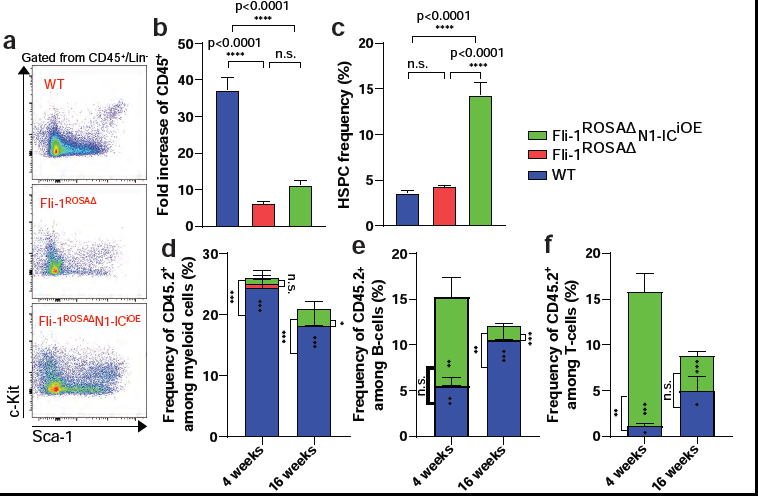
Enforcing Notch1 activation in Fli-1 deficient HSPCs restores regenerative expansion capacities and HSC function. WT, Fli-1^ROSAΔ^, and Fli-1^ROSAΔ^ with conditionally inducible Notch1 internal component overexpression transgene (Fli-1^ROSAΔ^N1-IC^iOE^) BM LSK HSPCs were isolated and expanded as described in Fig. S5A. Harvested cells were analyzed by flow cytometry and/or flow sorted for LSK HSPCs which were competitively co-transplanted with congenic SJL BM cells into lethally irradiated congenic SJL recipient mice. **a,** Representative flow dot plots for WT, Fli-1^ROSAΔ^, and Fli-1^ROSAΔ^N1-IC^iOE^ HSPCs post expansion, gated from the same number of CD45^+^/Lin^−^ cells. Note that Notch1 overexpression increased LSK frequency yet was not able to restore high c-Kit expression levels as WT LSK cells. **b,** Frequency of hematopoietic cells was determined by flow cytometry, number of cells was determined out of total MNC counts, and fold expansion was calculated. One-way ANOVA multiple comparisons was used; n=4 BM donor mice per genotype, with 2 technical replicates per donor. **c,** HSPC frequency as determined by flow cytometry. One-way ANOVA multiple comparisons was used; n=4 BM donor mice per genotype, with 2 technical replicates per donor. **d-f,** Frequency of chimerism among myeloid cells, B-cells, and T-cells as determined by flow cytometry. One-way ANOVA multiple comparisons was used; n=8 recipient mice per genotype.

**Extended Data Fig. 7:**
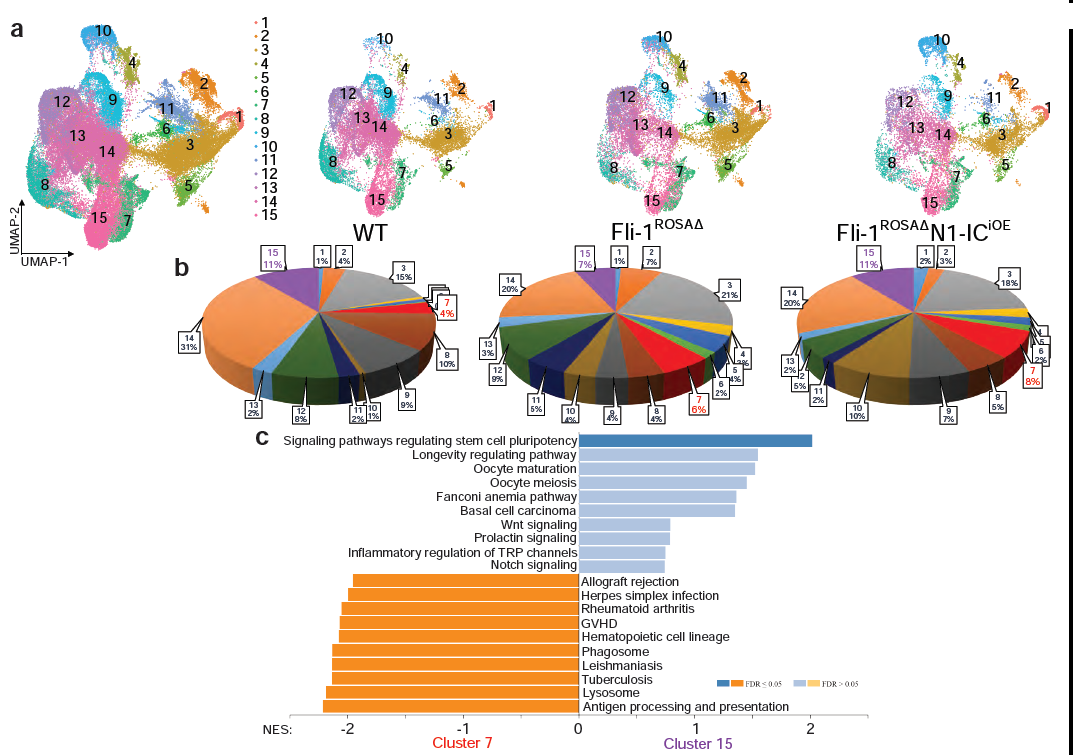
Enforcing Notch1 activation in Fli-1 deficient HSPCs restores the sub-population of transcriptionally defined activated HSCs. WT, Fli-1^ROSAΔ^, and Fli-1^ROSAΔ^ with conditionally inducible Notch1 internal component overexpression transgene (Fli-1^ROSAΔ^N1-IC^iOE^) BM LSK HSPCs were isolated and expanded. After 48 hours of expansion in co-culture, cells were harvested and hematopoietic LSK HSPCs were sorted and applied for single cell RNA-seq analysis; n=4 per genotype “pooled” together. **a.** Dimensionality reduction by UMAP of single cell transcriptomes from co-cultured and sorted LSK HSPCs per global (left plot) and per WT, Fli-1^ROSAΔ^, and Fli-1^ROSAΔ^N1-IC^iOE^ phenotype, as indicated. **b,** Pie chart displaying the relative proportion of each cluster per WT, Fli-1^ROSAΔ^, and Fli-1^ROSAΔ^N1-IC^iOE^ phenotype. Note HSC cluster 15 (purple) and HSC cluster 7 (red). **c,** Gene ontology (GO) categories of biological processes enrichment for differentially expressed genes in scRNA-seq analyses of HSC Cluster 15 vs. HSC Cluster 7, presented in a bar plot. Positively enrichment scored processes are enriched in HSC cluster 15 while negatively enrichment scored processes are enriched in HSC cluster 7.

**Extended Data Fig. 8:**
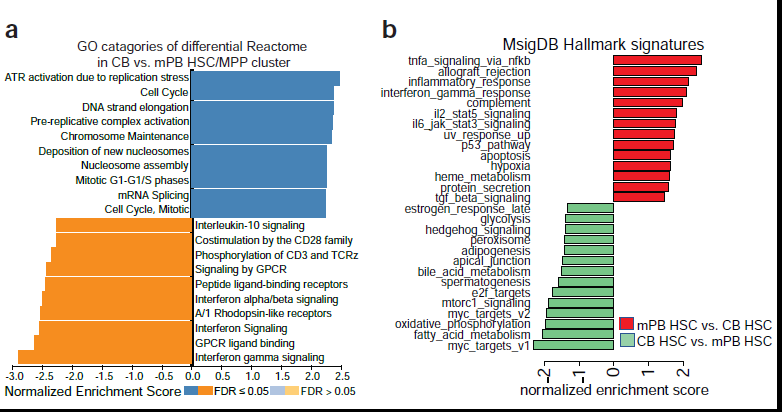
Transcriptional comparison distinguishes distinct reactome and hallmark signatures for neonatal and adult HSPCs. **a,** Gene ontology (GO) categories of reactome enrichment for differentially expressed genes in scRNA-seq analyses of cells from CB or mPB source in the HSC\MPP Cluster, presented in a bar plot. Positively enrichment scored processes are enriched in CB derived HSC\MPP cells while negatively enrichment scored processes are enriched in mPB derived HSC\MPP cells. **b,** Gene ontology (GO) categories of Hallmark signatures pathways enrichment for differentially expressed genes in RNA-seq analyses of sorted HSCs from CB or mPB source, presented in a bar plot. Positively enrichment scored processes are enriched in mPB HSCs while negatively enrichment scored processes are enriched in CB HSCs.

**Extended Data Fig. 9:**
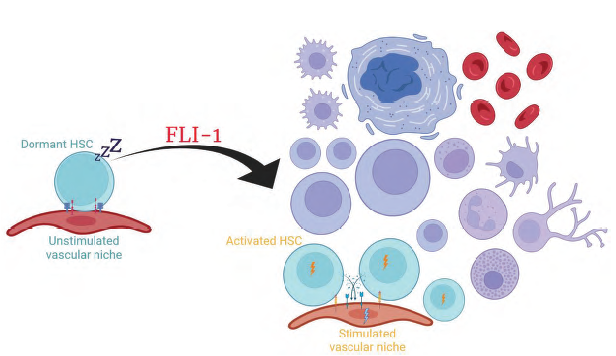
Schematic illustration of suggested model for FLI-1-driven active hematopoiesis from a quiescent HSC state. Attainment of FLI-1 transcriptional activity allows HSCs to adopt an active state. FLI-1 transcriptional activity steers HSCs out of their quiescence promoting HSPC expansion along with downstream differentiation to the essential blood cell lineages, enlarging the pool of functional HSCs capable of long-term engraftment and hematopoietic reconstitution post transplantation. Mechanistically, FLI-1 establishes the co-adoptability between an activated HSC to its supportive niche by mediating the stem cell-niche cell crosstalk and mutual sensing.

### Methods

#### Data reporting

No statistical methods were used to predetermine sample size. The investigators were not blinded to allocation during experiments and outcome assessment.

#### Animals

All animal experiments were performed under the approval of Weill Cornell Medicine Animal Care and Use Committee. Conditional mutants carrying *loxP*-flanked *Fli1* mice were generated by and obtained from Francois Morle’s laboratory^1^. These mice were backcrossed on a C57Bl/6J background for at least 10 generations. The following transgenic lines were applied in this study to induce specific Cre activity and gene inactivation/expression: Rosa26-Cre^ERT2^ mice were purchased from Jackson laboratory (stock #008463). TNR-eGFP were purchased from Jackson laboratory and have a C57BL/6J x SJL/J mixed background (stock #018322). Rosa26-N1-CD were purchased from Jackson laboratory (stock #008159). For human into mouse transplantation experiments NOD.Cg-*Kit^W-41J^Tyr*^+^*Prkdc^scid^Il2rg^tm1Wjl^* (NBSGW) were purchased from Jackson laboratory (stock #026622). For mouse into mouse transplantation experiments C57Bl/6J (CD45.2, stock #000664) and congenic B6.SJL-Ptprc^a^Pepc^b^/BoyJ (SJL, CD45.1, stock #002014) mice were purchased from Jackson laboratory. Mice carrying only a Cre transgene, or the additional indicated mutations (without a Cre transgene) were used as wild-type (WT) controls to exclude non-specific effects of Cre activation and/or of floxed alleles mutation. Gene deletion or activation was confirmed by qRT–PCR measurements from isolated LSK HSPCs, by inspection of RNA-seq data from isolated LSK HSPCs, and for some of the mutants also by observing eGFP expression. Male and female mice at 8–16 weeks of age were used for all experiments involving adult mice. All mouse offspring used for this study from all strains were routinely genotyped using standard PCR protocols. Sample size was limited by ethical considerations and background experience in stem cell transplantation (bone marrow transplantation) which exists in the laboratory for many years and other published manuscripts in the stem cell field, confirming a significant difference between means. No randomization or blinding was used to allocate experimental groups and no animals were excluded from analysis. All mutated or transgenic mouse strains had a C57BL/6 background unless otherwise indicated. For Cre^ERT2^ induction, Tamoxifen (Sigma, T5648-5G) was dissolved in sunflower oil (S5007-250ML) at the concentration of 20 mg/ml. Tamoxifen was intraperitoneally injected into mice at the dose of 100 mg/kg for 3 constitutive days.

#### Complete blood count (CBC) and mononuclear cell (MNC) count

Mice were anesthetized within anesthetic vaporizer delivery system, delivering isoflurane into a mouse holding chamber at constant flow rate of 2.5 of vaporized isoflurane (Henry Schein Animal Health 1169567762) at 3-5% for initial induction and then kept at 1-2% after the first 5 min with the oxygen adjustment (Tech Air, New York) to the flow rate of 1.0L/min. Blood was drawn from the retro-orbital plexus using capillary tubes (1.1mm X75 mm Color Code Red, Kimble Chase, 41B2501). One tube of blood (about 65 μl volume) was added into 195 μl of 10 mM EDTA/PBS buffer. Following brief votexing, samples were acquired using ADVIA120 hematology analyzer (Siemens). WBCs, RBCs, platelets, hemoglobin, hematocrit, as well as other differential blood parameters were obtained using this method. Blood, bone marrow, and cultured MNC cells were also counted using hematocytometer and EVOS imaging system (ThermoFisher Scientific) following dilution with Turk’s solution (EMD Millipore).

#### Transplantation assays

In all mouse into mouse transplantation experiments for engraftment purposes (non-homing experiments), mice were lethally irradiated with 475rad + 475rad doses with 5 hours interval in between, using an RS 2000 Biological Research X-ray Irradiator (Rad Source Technologies). Mice were transplanted with cells 24 hours post irradiation. For generation of chimeric mice, 2×10^5^ Lineage depleted BM cells from either WT or Fli-1^ROSAΔ^ mice were transplanted into C57Bl/6J WT recipient mice and hematopoietic BM recovery was allowed to proceed for at least 16 weeks before any other experimental procedure. For competitive transplantation assays testing HSC activity, 500 sorted (see flow and sorting section) LSK cells (per genotype and/or condition) were mixed with 4×10^5^ total BM cells from congenic SJL mice and transplanted into lethally irradiated WT congenic SJL mice. For generation of mixed BM chimeras, 2×10^5^ total BM cells from WT or Fli-1^ROSAΔ^ chimeric mice were mixed with 2×10^5^ total BM cells from congenic SJL mice and transplanted into lethally irradiated WT congenic SJL mice.

In human into mouse transplantation experiments for engraftment purposes, cultured cells were harvested, collected from each experimental well, and transplanted directly into BSGW immunodeficient mice without any myeloablative pre-conditioning. For secondary transplantation experiments, total bone marrow was harvested from primary recipients after 4 months of engraftment, incubated overnight in human expansion media and transplanted again into BSGW immunodeficient mice without any myeloablative pre-conditioning for additional 4 months.

To assess levels of chimerism, at indicated time points, mice were anesthetized (as previously described), blood was drawn (as previously described) or bone marrow and spleen were retrieved following recipient mouse sacrifice, RBCs were lysed for 10 min at 4°c using RBC lysis buffer (Biolegend), washed twice, stained for indicated flow markers prior, and acquired by flow cytometry (see flow and sorting section).

#### Methylcellulose assay (CFU-C assay)

A total of 20,000 bone marrow MNCs were seeded per 35 mm dish with MethoCult media (M3434, STEMCELL Technologies) or 100,000 bone marrow MNCs with MegaCult-C (04900 and 04902, STEMCELL Technologies) supplemented with murine Il-3 (10 ng/mL), human Il-6 (20 ng/mL), and murine TPO (50 ng/mL) (all from Peprotech) and cultured at 20% oxygen for 14 days. Megakaryocyte colonies were fixed and stained for acetylcholinesterase accordingly to manufacturer’s (STEMCELL Technologies) instructions. The type and number of colony-forming units were determined and scored. To help to visualize and define colony type, at day 14-post methylcellulose culture, CFUs were imaged using an automated colony counter, STEMvision (STEMCELL technologies).

#### Microscopic imaging

For bright field and fluorescent live-cell microscopy, a widefield setup was used. Live imaging was performed with a Zeiss Axio Observer Z.1 and a 10X/0.3NA objective with a reduced condenser aperture for enhanced contrast. Recordings were acquired using a sCMOS with 6.5µm2 pixels (Hamamatsu Flash4.0v2). Live experiments were performed within an incubation chamber at 37°c with 5% CO_2_, 5% O_2_ and high humidity (Zeiss Module S1 from Pecon). For acquisition of one-week movies of co-cultures, imaging acquisition intervals were set to 30 min. Acquisition of Hematoxylin and Eosin (H&E) stained bone marrow images was performed under the same setup using a 20X/0.8 objective paired with a Zeiss AxioCam 305 color camera. Harvested bones were fixed over-night in 4% PFA at 4°c, washed 3 times with PBS for 10 minutes each time. Bones were then decalcified for 48h using 100mM EDTA solution and washed 3 times with PBS for 10 minutes each time. Next, decalcified bones for paraffined histology were kept in 70% Et-OH and were sent to Histoserv, Inc. to be processed for: H&E. Data was analyzed using Zeiss Zen 2.6 software.

#### Intravital confocal and multiphoton microscopy for BM homing assays

Cell culture and transplantation: Lineage depleted HPCs were expanded for 4 days in culture with E4orf1-HUVEC niche cells, Fli-1 deletion was induced by introduction of 1 ng/mL 4-OHT, and cells were further cultured for 48 hours. After 6 days, cultures were harvested and LSK HSPCs were sorted. WT or Fli-1^ROSAΔ^ LSK HSPCs were stained with 10 µM DiI or DiD in hematopoietic expansion media respectively for 20 min at 37°c. A total of 8,000 collected Dil-Fli-1^ROSAΔ^ and 8000 DiD-WT cells in Ca^2+^/Mg^2+^-free phosphate-buffered saline (D-PBS) were then transplanted via retro-orbital injection into an anesthetized 8-12-week-old C57Bl/6J mice. Before transplantation, recipient mice were sub-lethally irradiated using an x-ray irradiator with a split dose of 300rad with a 5-hour interval between the two doses. Note: To control the dye labeling and detection efficiency, the DiI and DiD dye colors were swapped between WT and Fli-1^ROSAΔ^ LSK HSPCs in replicate experiments.

*In vivo* Imaging: Mice were anaesthetized with an induction dose of 3–4% isoflurane and a maintenance dose of 1.5–2% isoflurane. Mice were deemed anaesthetized by the toe pinch method. The hair on the calvarium was removed with a mechanical trimmer and then the skin was wiped with alcohol. The mice were then mounted in a designed heated mouse holder. Next, a calvarial skin flap was created with a U-shaped incision to reveal the underlying calvaria. A drop of D-PBS was applied to the skull as the immersion fluid. The mice were transferred to the stage of a multiphoton/confocal laser-scanning video-rate microscope and an Olympus 25×1.05 numerical aperture water-dipping objective was used for the imaging. The calvarial BM was imaged for 1-3 hours. Z-stack images were acquired with 2-3 µm step size. The excitation wavelength from an Insight X3 (Specra-Physics) was set at 1040 nm (two-photon) for DiD (Invitrogen), DiI (Invitrogen), and second harmonic generation (SHG) from collagen in the bone. The contrast and brightness of figure images and movies were adjusted for display purposes only. After imaging was completed, mice were dismounted from the stage and the skin flap was closed with 6-0 vinyl sutures (Ethicon) to enable repeat imaging the next day. Triple antibiotic ointment (bacitracin, neomycin, and polymyxin-B sulfate) was applied to the top of the surgical site to minimize the chance of infection. Mice were put in a heated cage and monitored until fully awake. Next, 24 hours after transplantation, mice were again prepared as described above and imaged to locate the homed LSK HSPCs in the BM. After locating the homed cells, mice were immediately injected with 100 µl of a vascular label, Fluorescein-Dextran (2,000,000da MW), retro-orbitally to visualize blood vessels (excited with the 1040 nm light). After imaging, mice were removed from the microscope and euthanatized.

#### Flow cytometry analysis and sorting

Cells were isolated and processed as described for each experimental procedure. Cells were incubated with mouse and/or human FcR block (Biolegend) in 50μl MACS buffer for 20 min at 4°c. Antibodies were added to resuspended cells as indicated per experiment (see antibody list) and cells with antibodies were incubated for 30 min at 4°c. For all the experiments, unstained sample or fluorescence-minus-one (FMO) rule was used for gating of positive populations. Unstained control and single-stained cells or ultracompensation beads (eBiosciences) were used to calculate and compensate for the fluorescence spillover from all the channels. Samples were acquired using a BD LSR II machine for flow cytometry or sorted using a BD ARIA 2 sorter machine. Dapi staining was performed at a final concentration of 1μg/ml prior to acquisition and/or sorting, to exclude dead cells. Data was analyzed using DIVA and FlowJo software. For apoptosis analysis, harvested and stained cells were washed in MACS buffer, and resuspended in Annexin V binding buffer (BD Pharmigen). AnnexinV was added accordingly to manufactures instructions. Dapi was added and samples were acquired on BD LSRII in less than 30 min post preparation. Prior to murine HSPCs sorting (and for other experimental procedure where mentioned), mature lineage positive hematopoietic cells were depleted using the direct lineage depletion kit (Miltenyi) accordingly to manufacturer’s instructions.

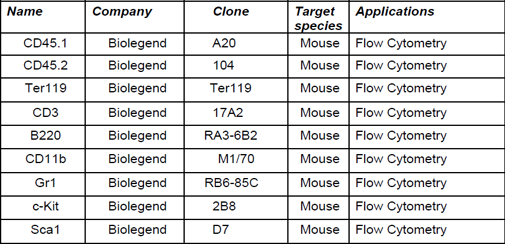
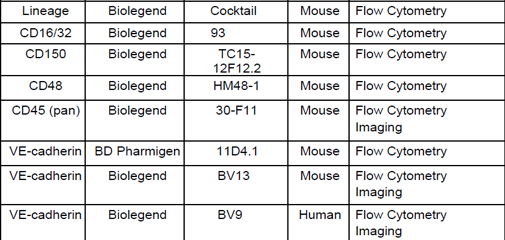
Antibodies table for flow and imaging.

#### Cell culture media

All in vitro cell preparations were routinely tested for mycoplasma contamination. Endothelial cell and co-culture harvesting were always performed using accutase solution (Corning).

*Endothelial cell media.* M199 (Hyclone), 20% FCS (Corning), 2mM glutagro supplement (Corning), Antibiotic Antimycotic solution (Corning), MEM nonessential amino acids (Corning), 20 mM HEPES (Invitrogen), 100 μg/mL heparin (Sigma), 25 μg/mL endothelial mitogen (Alfa Aesar) and 10 μM SB431542 (Tocris).

*Hematopoietic expansion media.* StemSpan SFEM (STEMCELL Technologies), 10% KnockOut Serum Replacement (Invitrogen), 50 ng/mL human/mouse c-Kit ligand (SCF, Peprotech), 50 ng/mL human/mouse TPO (Peprotech), and 50 ng/mL human/mouse Flt3l (Peprotech), 2mM glutagro supplement (Corning), Antibiotic Antimycotic solution (Corning), MEM nonessential amino acids (Corning), 20 mM HEPES (Invitrogen), and 100 μg/mL heparin (Sigma).

#### Generation of vascular niche cells

E4orf1-HUVECs that were used for HSPC expansion experiments and EHT conversion through reprogramming experiments, were produced as previously described^2^, with some modifications. Briefly, human umbilical vascular endothelial cells (HUVECs) were isolated as previously described^2^ and cultured in HUVEC media. The *E4orf1* gene from human adenovirus subtype 52 (GenBank accession No. ABK35065.1) was cloned into a pCCL-PGK lentivirus vector. HUVECs were transduced at 60-70% confluency with E4orf1 lentivectors at MOI of 1 supplemented with 4 ug of polybrene. HUVECs were incubated with E4orf1 lentivirus for 48 hours followed by 14 days selection using serum free X-VIVO medium (Lonza). E4orf1-HUVECs were used before reaching passage 20 in culture.

Generation of AGM-derived embryonic niche cells for embryonic EHT in vitro conversion experiments was previously described (AGM AKT-ECs)^3^. Briefly, AGM was isolated from e10.5 embryos, digested as described for EC isolation, and ECs were sorted, cultured and transfected with previously described^4^ myrAKT lentivector. Only lines selective for optimal EHT supportive capacity were applied for the EHT conversion experiments described in this manuscript.

#### Lentivirus production

Production of lentiviral delivery particles for E4orf1 was previously described in detai_l_^2^. Briefly, HEK 293T Lenti-X cell line (Clontech) cells were cultured in DMEM (Gibco) supplemented with 10% FCS (Corning), 2mM glutagro supplement (Corning), and Antibiotic Antimycotic solution (Corning) and transduced using Lenti-X packaging single shot (Clontech) following the manufacturer’s instructions (w/o antibiotics). Viral particles were concentrated using Lenti-X concentrator (Takara) and final titer was determined using Lenti-X p24 Rapid Titer Kit (Takara).

#### Construction of IVT templates and synthesis of modified-RNAs

Clean PCR products generated with plasmid templates purchased from GenScript were used as the template for mRNA. Modified-RNAs were generated by transcription *in vitro* with a customized ribonucleoside blend of ARCA; 30-O-Me-m7G (50) ppp(50)G (Trilink Biotechnologies); GTP; ATP; CTP (Life Technologies) and N1-methylpseudouridine-50-triphosphate (Trilink Biotech-nologies). The modified-RNA was purified with the MEGA clear kit (Life Technologies) according to the manufacturer’s instructions or using Amicon Ultra-4 Centrifugal Filter Unit 4 mL,10 kDa (Millipore Sigma) and treated with Antarctic Phosphatase (NEB). It was then re-purified with the MEGA clear kit. Then, modified-RNA was quantified using a Nano Drop spectrometer (Thermo Scientific), precipitated with ethanol and ammonium acetate, and re-suspended in 10 mM Tris-HCl and 1 mM EDTA at the final required concentration.

#### HSPC *in vitro* expansion s2tudies

Mouse hematopoietic BM sorted LSK HSPCs from different mouse strains were introduced into a co-culture with E4orf1-HUVEC niche cells (unless otherwise indicated), with hematopoietic expansion media, as previously described^5^, with few modifications. Deletion of Fli-1, reporter activation, or N1-CD overexpression, was induced by introduction of 1 ng/mL 4-OHT (Sigma-Aldrich). Hematopoietic expansion media was supplemented every other day post 4-OHT induction. Sorted BM LSK HSPCs were allowed to recover and enter the expansion phase for 48 hours prior to Fli-1 deletion. HSPCs were expanded for 6 days post Fli-1 deletion and/or other genomic induction and harvested for further flow sorting and other experimental procedures. For scRNAseq analysis HSPCs were isolated 48 hours post 4-OHT induction.

Human donor samples arrived from the Department of Pathology and were approved by the institutional review board at Will Cornell Medicine as part of the IRB protocol. Human hematopoietic CD34^+^ HSPCs from either cord blood (CB) or mobilized peripheral blood (mPB) donors, were isolated using human CD34 microbead kit (Miltenyi Biotec) and by passing twice through LS columns (Miltenyi Biotec) attached to a magnetic stand, achieving a purity >95%. Isolated Human hematopoietic CD34^+^ HSPCs were cryo-frozen. Cells were thawed prior to experimental procedures and were allowed to recover in human hematopoietic expansion media for few hours. For modified-RNA transduction, Neon electroporation transfection system (Invitrogen) was applied. For a suspension containing 1×10^5^ hematopoietic CD34^+^ HSPCs, 2 µg of GFP (control) or scrambled modified-RNA (control) or FLI-1 modified-RNA was introduced and electroporated accordingly to manufacture’s instruction using the following settings: 1700V, 10ms, 3 pulses. Immediately after electroporation cells were introduced into a co-culture with E4orf1-HUVEC niche cells with hematopoietic expansion media, at a ratio of 1:3 (HSPCs:E4orf1-HUVECs). Hematopoietic expansion media was supplemented every other day for 1 week. For scRNAseq analysis of CB and mPB HSPCs, cells were isolated and sorted 48 hours post introduction into co-culture with E4orf1-HUVEC niche cells.

#### Bulk RNA-seq

Mouse and human HSPCs were flow sorted as described. RNA was extracted and purified with Arcturus PicoPure RNA isolation kit (Applied Biosystems). RNA concentration and integrity were measured using Agilent 2100 Bioanalyzer (Agilent). RNA integrity was indicated by the RNA integrity number (RIN). RNA samples with sufficient concentration and RIN greater than 8.0 were further processed for poly-A selection and non-stranded cDNA library preparation using Truseq library preparation kit (Illumina). DNA library was then sequenced using Illumina HiSeq4000, PE read, 50 cycles. Raw fastq files were checked for quality with FastQC^6^ and adapters trimmed with Cutadapt^7^.

For mouse HSPC analysis, reads were aligned to mouse reference genome (mm10) using STAR aligner^8^. Raw gene counts were quantified using featureCounts from the Subread package^9^. StringTie2^10^ was used to quantify Transcripts Per Million (TPM) expression values. After further filtering and quality control, R package edgeR^11^ was used to calculate library size normalized FPKM and Log2 counts per million (CPM) values and perform differential gene expression analysis. FPKM normalized bigWig files were generated using deeptools^12^.

For human CB versus mPB HSPC comparison, data was normalized, and variance stabilized as a cohort using DESeq2 v1.18.1. Pathway analysis was performed using the gsva function of the GSVA_1.34.0 R package and GSEA_4.1.0 with 2000 permutations. For the GSEA analysis, a rank file was obtained by ranking all genes using the formula -log10(limma t value) that compared gene differential expression between the mPB and CB groups. GSVA results were visualized using the R gplots_3.1.1 heatmap.2 and ggplot2_3.3.3 boxplots.

#### Bulk ATAC-seq

We have followed the previously described Omni-ATAC-seq protoco_l_^13^. Briefly, LSK HSPCs were flow sorted as described. Cells were spun down for at 500rcf for 5 min at 4°c in a fixed angle. Next, cells were resuspended in RSB buffer and pipetted 3 times, followed up by 3 min incubation on ice. Cell lysate was washed, and nuclei pellet was acquired after spin at 500rcf for 10 min at 4°c. Nuclei pellet was resuspended in transposition mixture at incubated at 37°c for 30 min in a thermomixer with 1000 rpm mixing speed. Cleanup reaction was performed with a Zymo DNA clean, and concentrator-5 kit (Zymo). DNA was eluted and amplified using NEBNext X2 MasterMix (NEB). Following additional amplification and cleanup, library concentration was determined using the KAPA library quantification kit (KAPA). DNA library was sequenced on Illumina, using HiSeq4000, PE read, 50 cycles. Raw fastq files were checked for quality with FastQC^6^ and adapters trimmed with Cutadapt^7^. Reads were aligned to the mouse (mm10) reference genome using Bowtie2^14^ with parameters ‘--very-sensitive -X 2000’. Mitochondrial, duplicate, and low-quality reads were removed with only properly paired reads retained. Reads were shifted + 4bp and - 5bp on the positive and negative strands respectively. Peaks were called using MACS2^15^ with parameters ‘-f BAMPE -g 1.87e9 --keep-dup all –nomodel --nolambda --call-summits’. BigWig files were generated using deeptools^12^ normalized by reads per genomic content (1x normalization). Differential accessibility was performed with DESeq2^16^. Computational footprinting was performed with the HINT-ATAC^17^ and TOBIAS^18^ frameworks using the HOCOMOCOv11^19^ and JASPAR 2020^20^ vertebrate core transcription factor motif database respectively. For footprinting, biological replicate alignments were merged and PCR duplicate reads were removed with the Picard Toolkit^21^. Peaks were recalled on the merged alignment files with the same parameters.

#### Bulk ChIP-seq for H3K27ac

Mouse LSK HSPCs were flow sorted as described. Nuclei were extracted using NP40 buffer and digested into single nucleosomes using diluted MNase (NEB), followed by O/N incubation with the H3K27ac (Active Motif) antibody. The nucleosomes were washed twice with low salt wash buffer and twice with high salt wash buffer at 4°c in a rotator. DNA was eluted using SDS/NaHCO3 buffer at 65°c for 2 hours, extracted using phenol:chloroform:isoamyl alcohol (Invitrogen) and precipitated using 2-isopropanol (Sigma) at -80°c for O/N or longer durations. DNA library was prepared using Kapa Hyper Kit (KAPA) according to manufacturer’s instructions. Adaptors were from KAPA Single-Indexed Adaptor Kit (KAPA). DNA was purified using the DNA purification kit AMPure XP (Beckman Coulter). DNA library was sequenced on Illumina, using Hiseq 4000, single read, 50 cycles. Raw fastq files were checked for quality with FastQC^6^ and adapters trimmed with Cutadapt^7^. Reads were aligned to the mouse (mm10) reference genome using Bowtie2^14^ with parameters ‘--very-sensitive -X 2000’. Mitochondrial, duplicate, and low-quality reads were removed. Peaks were called using MACS2^15^ using input controls with parameters ‘-f BAM -g 1.87e9 --keep-dup all’ BigWig files were generated using deeptools^12^ normalized by reads per genomic content. Differential peak analysis for regulation of histone acetylation was performed with DESeq2^16^.

#### Differential peak motif enrichment analysis (MEA)

Peaks with an FDR < 0.05 were selected from the ATAC-seq and H3K27ac ChIP-seq comparisons of knockout to wildtype using the above comparison method. Peaks in each of the two assays were then analyzed for enriched KO and WT motifs using the HOMER software’s findMotifsGenome function^22^ with -size given and the default motif database.

#### Generation of *RNA vs. ATAC* dot plots

Mouse HSPC differentially expressed genes between knockout and wild-type with an FDR value less 0.05 and absolute log fold change value greater than 2.5 were selected for further interrogation. Differentially accessible regions from the ATAC-seq were annotated using the ChIPseeker R library^23^ and only included if this fell within 500bp up or downstream of a transcriptional start site. Accessible regions were then only included if they had a FDR < 0.05, and an absolute log fold change value greater than 0.75 in comparing knockout to wild-type. For any gene annotated to multiple peaks, the greatest absolute value log fold change was selected for plotting to avert one to many relationships and visualize the chromatin changes with the largest magnitude.

#### Integrative Genomics Viewer (IGV) 2.5.3 was used to display peaks tracks for RNA, ATAC (including footprinting), and ChIP (set to mouse mm10 genome)

##### Gene enrichment analysis

Gene enrichment for gene ontology (GO) biological processes was performed using WebGestalt^24^. Gene set enrichment analysis (GSEA) for selected pathways was performed using the Broad Institute and UC San Diego software^25, 26^ ranked by log fold change.

#### Single cell/nuclei RNA/ATAC-seq analyses

##### Multiome single nuclei RNA/ATAC-seq analysis

Mice were sacrificed, bones were harvested, flushed, crushed, and processed to flow sort SLAM LSK HSPCs as described above. SLAM LSK HSPCs from 20 C57BL/6 mice were pooled together, washed, and resuspended in 0.04% BSA/PBS solution. Sorted cell suspension was further processed for nuclei isolation with the Chromium Next GEM Single Cell Mutiome Reagent Kit (10x Genomics, product code # CG000365 and 1000338) using 10X Genomics’ Chromium Controller. Briefly, 200K sorted SLAM LSK cells were suspended in Lysis Buffer and incubated on ice for 5 min. Chilled Wash Buffer was added, mixed well, and cells were centrifuged at 500 rcf for 5 min, then washed twice and resuspended in diluted Nuclei Buffer. The nuclei concentration was determined by Bio-Rad TC20 Cell Counter, and immediately proceeded to Chromium Next GEM Single Cell Mutiome ATAC + Gene Expression. Nuclei suspensions were incubated in a Transposition Mix that includes a Transposase, which preferentially fragmented the DNA in open regions of the chromatin. Simultaneously, adapter sequences were added to the ends of the DNA fragments. Single Cell Multiome ATAC + GEX Gel Beads included a poly(dT) sequence that enabled the production of barcoded, full-length cDNA from poly-adenylated mRNA for gene expression (GEX) library and a Spacer sequence that enabled barcode attachment to transposed DNA fragments for ATAC library. Transposed nuclei were loaded into Chromium Controller to generate GEMs by combining barcoded Gel Beads, transposed nuclei, a Master Mix, and Partitioning Oil on a Chromium Next GEM Chip J. Upon GEM generation, the Gel Bead was dissolved. Oligonucleotides containing an Illumina P5 sequence, a 16 nt 10x Barcode (for ATAC), and a Spacer sequence were released. In the same partition, primers containing an Illumina TruSeq Read 1 (read 1 sequencing primer), 16 nt 10x Barcode (for GEX), 12 nt unique molecular identifier (UMI), and a 30 nt poly(dT) sequence were also released. The primers were mixed with the nuclei lysate containing transposed DNA fragments, mRNA, and Master Mix, that included reverse transcription (RT) reagents. Incubation of the GEMs produced 10x Barcoded DNA from the transposed DNA (for ATAC) and 10x Barcoded, full-length cDNA from poly-adenylated mRNA (for GEX). This was followed by a quenching step that stopped the reaction. GEMs were broken and pooled fractions were recovered. Silane magnetic beads were used to purify the cell barcoded products from the post GEM-RT reaction mixture, which included leftover biochemical reagents and primers. Barcoded transposed DNA and barcoded full-length cDNA from poly-adenylated mRNA were amplified via PCR to fill gaps and for generating sufficient mass for library construction. The pre-amplified product was used as input for both ATAC library construction and cDNA amplification for gene expression library construction. P7 and a sample index were added to pre-amplified transposed DNA during ATAC library construction via PCR. The final ATAC libraries contained the P5 and P7 sequences used in Illumina bridge amplification. Barcoded, full-length pre-amplified cDNA was amplified via PCR to generate sufficient mass for gene expression library construction. Enzymatic fragmentation and size selection were used to optimize the cDNA amplicon size. P5, P7, i7 and i5 sample indexes, and TruSeq Read 2 (read 2 primer sequence) were added via End Repair, A-tailing, Adaptor Ligation, and PCR. The final gene expression libraries contained the P5 and P7 primers used in Illumina bridge amplification. Sequencing these libraries with Illumina NovaSeq6000 using 100 cycles kits produced a standard Illumina BCL data output folder that included paired end Read 1N and Read 2N used for sequencing the DNA insert, along with the 8 bp sample index in the i7 read and 16 bp 10x Barcode sequence in the i5 read (50+8+24+49). Chromium Single Cell Multiome Gene Expression libraries comprised cDNA insert with standard Illumina paired-end constructs which began with P5 and end with P7. Sequencing these libraries produced a standard Illumina BCL data output folder. TruSeq Read 1 was used to sequence 16 bp 10x Barcodes and 12 bp UMI, while 10 bp i5 and i7 sample index sequences were the sample index reads. TruSeq Read 2 was used to sequence the insert (28+10+10+90).

Next, to remove ambient RNA, random UMI swapping, and technical artifacts, CellBender^27^ was run on the single-nuclei RNAseq raw Cell Ranger (10X Genomics) feature counts with 150 epochs, 0.01 fpr, and the total droplets/expected cells selected based on quality control web summary elbow plot. DoubletDetection^28^ was also run on the raw features and the overlapping barcodes filtered from the Cellbender expression results. To further filter doublets and poor quality cells, cells with greater than 250 or less than 3400 genes, greater than 14% mitochondrial genes, or RNA counts greater than 350 or less than 10,000 were filtered. Ribosomal and mitochondrial RNA counts were excluded from downstream analysis. Cells with fewer than 2,740 or greater than 69,000 ATAC counts were further excluded. The alignment gene transfer format annotation file was used to define transcriptional start site (TSS) regions as ±3 kb from TSS. TSS enrichment scores were computed using the Signac TSSErichment function. Cells with TSS enrichment <3 were discarded. Peaks were recalled using MACS2^15^ through the Signac CallPeaks function, parameters effective.genome.size = 1.87e9, extsize = 150, shift = 75, -nomodel, --call-summits --nolambda --keep-dup all. Non-standard chromosomes were removed along with any mitochondrial or ChrY chromosome peaks. Regions that overlapped with the mm10 blacklist region were further removed. Peaks were centered on the summit and extended 250bp in each direction. Peaks were sorted by score and any overlapping peaks had the peak with the largest score included. The subsequent top 100,000 peaks sorted by greatest score were included for further analysis. Gene expression was normalized using sctransform^29^ version ‘v2’ prior to running principal component analysis. Peaks/accessible regions were processed using the Signac methods RunTFIDF, FindTopFeatures (with min.cutoff = ‘q0’) and RunSVD. Multimodal neighbors were found by constructing a weighted nearest neighbor graph^30^ using ^31^ the Seurat FindMultiModalNeighbors function. The first 50 principal components from the expression data and the 2^nd^ through 50^th^ latent semantic index (LSI) components from the reduced ATAC data were used. The SLM algorithm was then applied to find clusters with a resolution of 0.3. HSC, MPP, EryP, and MkP clusters were annotated based on RNA markers identification using Seurat FindAllMarkers. FindMarkers Seurat function with the Wilcoxon Rank-Sum test identified differentially expressed genes. Differential accessible regions were found using the FindMarkers Seurat function with the logistic regression framework with the number of peaks as a latent variable. Differential motifs were found using the FindMarkers Seurat function with Wilcoxon Rank-Sum test with the average difference calculation on the chromVAR z-score (rowMean). Motif analysis was performed using chromVAR with the CIS-BP 2021 curated mouse motif database database^32^. Cell cycle scores were calculated using the Seurat “CellCycleScoring” function. The HSC signature was calculated using the Seurat “AddModuleScore” function with default parameters with the Rodriguez et al. HSC1 gene list^33^. The Nebulosa R package^34^ was used to clearly visualize gene signatures by mitigating sparsity through kernel density estimation. UMAPs and other figures were designed using SCpubR^35^. Heatmaps for RNA and Chromvar motifs activity present the top 30 genes and motifs per cell type by avg_log2FC (RNA) and fold.enrichment (motif), after selected by statistical significance of p adjusted value <0.05. Differential RNA and motif per cluster were determined by applying FindMarkers (RNA) and FindMotifs (motif) Seurat functions by comparing to nearest cell types. HSC and MPP cells were subset without EryP/MkP (n=2965) and had peaks re-called using the above methods. The same dimensional reduction and clustering approaches were re-applied using the first 35 principal comonents, the 2^nd^ through 40^th^ LSI components and a cluster resolution of 0.5 due to the smaller amount of cells. We removed a small number of HSC and MPP outlier cells (n=17), that were outliers in low dimensional space and had enrichment for erythroid cell markers (such as hemoglobin and etc.), and reclustered with the same HSC/MPP parameters. SCPubR was used to generate dot plot and violin plots of Chromvar motifs scores for selected TFs. For statistical analysis between HSC1 and HSC2 motifs scores presented on violin plots, Wilcoxon Rank Sum Test (rstatix library v0.7.0) was used to calculate P values as indicated.

##### General single cell RNA-seq procedures

Sorted cells were immediately submitted for 10X Chromium single-cell analysis. The single-cell suspension was loaded onto a well on a 10X Chromium Single Cell instrument (10X Genomics). Barcoding and cDNA synthesis were performed according to the manufacturer’s instructions. In brief, the 10X GemCode Technology partitions thousands of cells into nanolitre-scale gel bead-in-emulsions (GEMs), in which all the cDNA generated from an individual cell share a common 10X barcode. To identify the PCR duplicates, a unique molecular identifier (UMI) was also added. The GEMs were incubated with enzymes to produce full length cDNA, which was then amplified by PCR to generate enough quantity for library construction. Qualitative analysis was performed using the Agilent Bioanalyzer High Sensitivity assay. The cDNA libraries were constructed using the 10X Chromium single-cell 3′ Library Kit according to the manufacturer’s original protocol. In brief, after the cDNA amplification, enzymatic fragmentation and size selection were performed using SPRI select reagent (Beckman Coulter) to optimize the cDNA size. P5, P7, a sample index and read 2 (R2) primer sequence were added by end repair, A-tailing, adaptor ligation and sample-index PCR. The final single-cell 3′ library contains standard Illumina paired-end constructs (P5 and P7), Read 1 (R1) primer sequence, 16-bp 10X barcode, 10-bp randomer, 98-bp cDNA fragments, R2 primer sequence and 8-bp sample index. For quality control after library construction, 1 μl of the sample was diluted 1:10 and run on the Agilent Bioanalyzer High Sensitivity chip for qualitative analysis. For quantification, the Illumina Library Quantification Kit (KAPA) was used. DNA libraries were sequenced on Illumina, using NovaSeq6000, PE read, 50 cycles.

The Cell Ranger pipeline (10X Genomics) was used to align reads to the mm10 (for murine samples) or to the GRCh38 (for human samples) reference genomes and generate feature barcode matrices.

##### Single cell RNA-seq analysis of engrafted murine HSPCs

Bone marrow was retrieved from recipient mice >16 weeks post transplantation, and viable donor immature hematopoietic population enriched for HSPCs (CD45.2^+^Lin^−^ Dapi^−^) was flow sorted. For each donor genotype BM samples were pooled from 5 recipient mice.

The Monocle 3 (v.3_0.2.3.0) platform was used for downstream analysis^36^, combining data for cells from each sample for downstream analysis, using a negative binomial model of distribution with fixed variance, normalizing expression matrices by size factors. Counts for UMI (unique molecular identifiers) and unique genes expressed per cell are shown in boxplots in supplementary figures for each sample (showing median values and interquartile ranges; upper/lower whiskers show 1.5X interquartile range with outliers shown as individual dots). Low quality cells were excluded using cut-offs for UMI per cell (<5000) and genes per cell (<1000). R scripts used for analysis are available upon request (R version 3.6.1). The preprocess_cds function was used to project the data onto the top principal components (excluding principal components that contributed little to the overall variance, num_dim=9). The align_cds function was used to remove batch effects between samples, using a “mutual nearest neighbor” algorithm, Batchelor (v.1.2.4)^37^. Uniform Manifold Approximation (UMAP)^38^ was used for dimensionality reduction with the reduce_dimension function, using default settings. Clustering was performed by Leiden method with the cluster_cells function, with resolution=2E-3. Cell type classification was performed using the Garnette package (v.0.2.15)^39^ within Monocle 3. Marker genes based on established cell type-specific genes, as indicated below, were used to train a classifier data set (using the train_cell_classifier function with default settings), and classify cell types (using the classify_cells function, with cluster_extend set to TRUE in order to expand cell classifications to cells in the same cluster; settings cluster_extend_max_frac_unknown = 0.95 and cluster_extend_max_frac_incorrect = 0.45). Cluster-extend cell types were used for downstream analysis comparing samples.

Markers for cell type classification.

Markers for cell type classification
| Cell Type | Genes Expressed | Genes not expressed |
| --- | --- | --- |
| HSC | *Hoxb5, Fgd5, Cdkn1c, Procr, Slmaf1* | *Cd48, Flt3, Cd34* |
| MPP | *Cd34, Hlf* | *Hoxb5, Cdkn1c, Fgd5, Cebpe, Fcnb, Elane, Klf1, Gp1bb, Gata1, Csfr1, Siglech, Fcer1a, Il2rb, Procr, Vwf, Dntt, Rag2* |
| LPP | *Dntt, Rag2* | *Hlf, Procr, Gata2, Siglech, Ccl5, Il2rb, Fcer1a, Ms4a2* |
| MEP | *Gata1, Zfp1* | *Fcer1a, Ms4a2, Procr, Fgd5, Hoxb5, Ly6a, Flt3, Ccl5, Il2rb* |
| Monocyte | *Adgre1, Csf1r* | *Siglech, Fcnb* |
| Neutrophil | *Fcnb, Elane, Cebpe* | *Adgre1, Ly86, Siglech* |
| Dendritic | *Siglech* | *Adgre1, Fcnb, Elane* |
| Mast-Basophil | *Fcer1a, Ms4a2* | *Pf4, Klf1* |
| Lymphocyte | *Ccl5, Il2rb* | *Siglech, Csf1r* |

Gene-set scores were calculated for each single cell as the log-transformed sum of the size factor-normalized expression for each gene in signature/marker gene sets, as indicated, including two published HSC-specific gene sets: 29 genes representing a molecular overlap signature of adult murine bone marrow-derived HSC isolated by different index sorting strategies^40^ and 541 genes representing the molecular signature of serially-repopulating HSC by single cell clonal tracking^33^. Violin plots comparing gene-set scores between samples were generated using ggplot2 (v3.3.2) function with geom_violin() and geom_boxplot() (boxplots show median values and interquartile ranges; upper/lower whiskers show 1.5X interquartile range). For statistical analysis between samples, Wilcoxon Rank Sum Test (ggupbr package v0.4.0) was used to calculate P values as indicated. Differential gene expression was performed using regression analysis with the fit_models() and coefficient_table() functions in Monocle 3, to identify genes that were differentially expressed based on significance values (q value) adjusted for multiple hypothesis testing using the Benjamini and Hochberg correction method. Hematopoietic stem/progenitor cell types (HSC, MPP, and LPP) were compared between samples for this analysis, to identify differentially expressed genes in the whole genome (genome-wide) or using signature/marker genes sets that were utilized for gene-set scores as described above.

##### Single cell RNA-seq analysis of murine HSPCs from co-cultures

Sorted BM LSK HSPCs, from all the different transgenic mouse lines, were introduced into co-culture with vascular niche cells as described above. Two days (48 hours) post tamoxifen introduction into co-cultures containing expanding HSPCs, cultures were terminated, harvested, and LSK HSPCs were resorted and further processed for 10X chromium scRNA-seq analysis. Co-culture sorted LSK HSPC samples were loaded into Seurat. For each sample, cells without 100 features and features detected in less than 10 cells were discarded. Each sample was filtered individually for low quality cells and doublets. The WT sample had cells discarded with less than 350 or more than 3750 features and greater than 25% mitochondrial content. The Fli-1^ROSAΔ^ sample had cells discarded with less than 320 or more than 2000 features and greater than 15% mitochondrial content. The Fli-1^ROSAΔ^N1-IC^iOE^ sample had cells discarded with less than 325 features or more than 4200 features and greater than 35% mitochondrial content. The N1-IC^iOE^ sample had cells discarded if they had less than 350 or more than 4500 features and greater than 35% mitochondrial content. Each sample was normalized using sctransform and integrated using 3000 integration features and default parameters. Cell cycles were classified using the CellCycleScoring function along with the Seurat list of cell cycle genes. Principal component analysis was run, and the first 40 principal components were selected for finding neighbors and computed the UMAP based on the elbow method as well as the derivative of an interpolated function fit to the elbow plot showing little change for additional components. The FindClusters function was run with a resolution of 0.6 and HSC clusters were annotated based on the described above. Cells from HSC defined clusters #7 and #15 were selected for further interrogation and the FindMarkers function was used with the RNA assay and default parameters for differential expression analysis between the two populations and the different genetic conditions within each cluster. Gene set enrichment analysis was performed using WebGestalt.

##### Single cell RNA-seq analysis of human CB and mPB HSPCs

CB and mPB enriched CD34^+^ HSPCs were introduced to co-cultures with vascular niche cells, as described above, for 48 hours. CB and mPB CD34^+^ HSPCs were sorted from harvested cultures and further processed for 10X chromium scRNA-seq analysis. Both CB and mPB HSPC samples were first filtered for cells that had less than 100 features and features that occurred in less than 3 cells. To filter out low quality cells and doublets, CB HSPCs with less than 200 features or greater than 7200 features and greater than 12 percent mitochondrial content were discarded. Additionally, mPB HSPCs with less than 1000 features or greater than 6100 features and a percent mitochondrial content greater than 11 percent were removed. After merging the samples, the Seurat functions NormalizeData and ScaleData were run with default parameters. The top 4000 variable features were then identified using the FindVariableFeatures routine with the ‘vst’ selection method and cell cycles scored with CellCycleScoring method using Seurat’s list of cell cycle genes. Mitochondrial percentage and the cell cycle difference, calculated as the S.Score minus G2M.Score, were regressed out, the latter so the signal between cycling and non-cycling HSPCs was maintained. After running PCA reduction, Harmony^41^ was used to integrate the two different CB and mPB batches. Nearest neighbors and UMAP dimension reduction were calculated using the first 40 corrected Harmony embeddings. Additionally, Nearest neighbors were calculated with k=30 for k-nearest neighbor. The FindClusters function was run with a resolution of 2. To annotate cell types to clusters, a combination of known genes, markers found with the FindAllMarkers function (using the RNA assay with only positive markers and a p-adjusted value < 0.05). Cells were first annotated through the Azimuth reference-based mapping pipeline on human bone marrow^30^. Clustering and cluster cell assignment was finalized using known cell markers. Additional differential gene expression was performed between cell types, samples, and clusters using the FindMarkers Seurat function with an average log fold change limit threshold of 0.15 for genes between the two comparison groups. Gene set enrichment analysis was performed using WebGestalt.

##### Single cell ATAC-seq HSPC analysis of quiescent, active, and FLI-1 HSPC signatures

Publicly available data for human HSPC scATA-seq (EGAS00001004740) was analyzed, where cells from three sorted CD34^+^CD38^−^CD45RA^−^ and three CD34^+^CD38^+^ populations were processed on the 10X Genomics single cell ATAC-seq platform and retained based on default cellranger QC criterion and chromVAR depth filtering. Cellranger-reanalyze (1.1, 10x Genomics) was subsequently used to map reads over the sites identified in the publicly available human HSPC bulk ATAC-seq catalog (EGAS00001004742), and read counts were binarized. chromVAR (with default settings) was used to calculate the enrichment of the quiescent and active hematopoietic signatures identified in Takayama et al^42^, per each single cell, as well as for the HSPC FLI-1 CHIP-Seq^43^ peaks signature.

##### Statistical analysis

All statistical analyses were conducted with Prism (* *P* < 0.05, ** *P* < 0.01, *** *P* < 0.001, **** *P* < 0.0001; NS, not significant). All data are expressed as mean ± standard error (s.e.m) and all *n* numbers represent biological repeats, unless indicated otherwise. Unless indicated otherwise in figure legends, a student’s two-tailed unpaired *t*-test was used to determine the significance of the difference between means of two groups. One-way ANOVA or two-way ANOVA was used to compare means among three or more independent groups. Bonferroni post-hoc tests were used to compare all pairs of treatment groups when the overall *P* value was < 0.05. A normal distribution of the data was tested using the Kolmogorov–Smirnov test if the sample size allowed. If normal-distribution or equal-variance assumptions were not valid, statistical significance was evaluated using the Mann–Whitney test and the Wilcoxon signed rank test.

##### Data and code availability

All codes and mathematical algorithms applied in this manuscript will be provided upon request from the corresponding authors.

