## Supplementary figures and images for "Transcriptional Activation of Regenerative Hematopoiesis via Vascular Niche Sensing"

### Supplemental Table 2

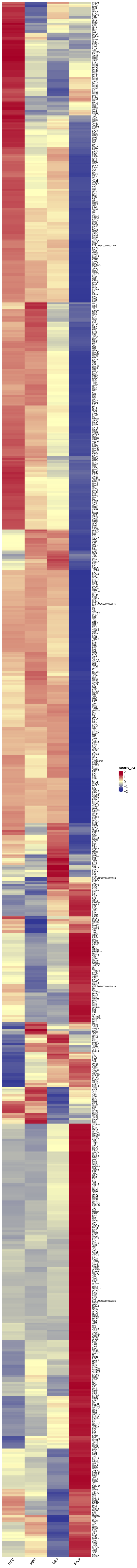
